# Caspase-8-mediated CYLD Cleavage boosts LPS-induced Endotoxic Shock

**DOI:** 10.1101/2024.04.03.587858

**Authors:** Jianling Liu, Ming Li, Mingyan Xing, Han Liu, Xiaoxia Wu, Lingxia Wang, XiaoMing Li, Xiaoming Zhao, Yangjing Ou, Yue Zhang, Fang Li, Qun Xie, Yongchang Tan, YangYang Wang, YangYang Xie, Hanwen Zhang, Hong Li, Yu Li, Yan Luo, Haibing Zhang

## Abstract

Caspase-8, a pivotal protease intricately involved in various cellular signaling pathways related to cell death and inflammation, has been identified as a contributor to cytokine production during septic shock. However, the mechanisms governing this regulatory role remain enigmatic. In this study, we uncovered that mice harboring a specific mutation in CYLD at its D215 position (*Cyld^D^*^215^*^A/D^*^215^*^A^* mutant mice), rendering CYLD resistant to caspase8 cleavage, exhibited marked protection against lethal endotoxic shock. Moreover, the removal of *Cyld* in *Caspase8^-/-^Mlkl^-/-^* mice restored their sensitivity to endotoxic shock, indicating Caspase8 promotes LPS-induced endotoxic shock by cleaving its substrate CYLD and maintaining CYLD stability confers resistance to endotoxic shock. Mechanistically, CYLD was found to catalyze the removal of LUBAC-mediated M1-linked ubiquitination of NF-kB p65 at K301/K303, thereby suppressing the nuclear translocation and activation of p65 for subsequent cytokines production. Notably, the cleaved N-terminal fragment of CYLD (named CP25) is secreted in a TRIF/Caspase8-dependent manner. Moreover, CP25 can be detected in the serum of septic mice, and its levels show a strong correlation with corresponding serum IL-1β, suggesting its potential utility as an inflammatory biomarker. Overall, these findings highlight the significance of CYLD cleavage as a promising therapeutic target and diagnostic marker for endotoxic shock.

## Introduction

Sepsis is a systemic inflammatory response syndrome triggered by an infection that leads to severe organ dysfunction as a result of an uncontrolled host responses^1,2^. Despite extensive research on its pathogenesis, sepsis continues to be the main cause of infection-related mortality wordwilde^1,3^. Lipopolysaccharide (LPS), a component derived from gram-negative (GN) bacteria, is a significant contributor to sepsis and plays a pivotal role in inducing inflammation^2^. LPS activates signaling through canonical and noncanonical inflammasome mediated pyroptosis^4,5^. Caspase1, activated by dimerization and auto-processing, controls maturation and secretion of IL-1β and IL-18 and induces pyroptosis^5–8^. In contrast, Caspase11 (Caspase4 and Caspase5 in humans) is directly activated by sensing cytosolic LPS and cleaves gasdermin-D (GSDMD) to form pore-forming GSDMD N-terminus fragment (GSDMD-NT), which allows the indirect activation of the noncanonical NLRP3 inflammasome and Caspase1 to promote activation of IL-1β and IL-18^4,9–11^.

Importantly, several studies have demonstrated that *Caspase8^-/-^Mlkl^-/-^* (or *Caspase8^-/-^Ripk3^-/-^*) mice, or those deficient in its adaptor protein FADD, exhibit increased resistance to endotoxic shock induced by LPS or bacterial infection^12–15^. These findings indicate that Caspase8 (or FADD), rather than RIPK3/MLKL, serves as the principal mediator of protective resistance^12–15^. However, the precise underlying mechanism responsible for this resistance remains unclear. Caspase8 functions in a regulatory role by cleaving various substrates such as cysteine proteases^16,17^. Notably, a recent study showed the involvement of Caspase8 in regulating innate immune responses through N4BP1 cleavage^18^. Another study demonstrated its role in facilitating TNF-induced necroptosis through the cleavage of CYLD at Asp215^19,20^. However, whether Caspase8 is involved in regulating the resistance to LPS-induced shock exhibited by *Caspase8^-/-^Mlkl^-/-^*(or *Caspase8^-/-^Ripk3^-/-^*) mice (or its adaptor FADD-deficienct) through CYLD cleavage remains unknown.

Ubiquitination is a post-translational modification that plays a crucial role in the involvement of various human diseases^21^. The linear ubiquitin chain assembly complex (LUBAC), composed of Heme-oxidized IRP2 ubiquitin ligase-1L (HOIL-1L), HOIL-1L-interacting protein (HOIP), and Shank-associated RH domain interactor (SHARPIN) subunits, connects linear ubiquitin chains to substrates^22,23^. CYLD was first identified as a tumor suppressor mutated in familial cylindromatosis^24,25^. Later studies have reported that CYLD is a deubiquitinating enzyme (DUB) that removes ubiquitin chains from substrates^26–29^. However, the role of CYLD in sepsis remains unexplored.

In this study, we demonstrated that Caspase8-mediated cleavage of CYLD at D215 plays a critical role in LPS-induced endotoxic shock. Our data revealed that *Cyld^D^*^215^*^A/D^*^215^*^A^* mice, expressing the D215A mutant resistant to Caspase8 cleavage, exhibited remarkable resistance to endotoxic shock, suggesting that non-cleaved CYLD prevented its degradation, thereby providing protection against LPS-induced endotoxic shock. Moreover, the deletion of *Cyld* in *Caspase8^-/-^Mlkl^-/-^* mice substantially restored their sensitivity to endotoxic shock compared to that in *Caspase8^-/-^Mlkl^-/-^* mice. This result further confirmed that maintaining CYLD stability was the key factor responsible for the resistance observed in *Caspase8^-/-^Mlkl^-/-^* mice. Mechanistically, CYLD, which functions as a deubiquitinase, directly interacts with P65 to regulate the removal of M1-linked ubiquitin chains from P65. Notably, CYLD, in collaboration with the LUBAC complex, exerts specific control over the balance of M1-linked ubiquitin on the K301/303 residues of P65, downstream of its detachment from IκBα. Furthermore, we observed that the cleaved N-terminal fragment of CYLD, named CP25, which lacks a signal peptide, could be secreted from the cytosol through a novel secretory pathway that depends on the TRIF/Caspase8 complex. The release of CP25 was detected in serum, highlighting its potential as a biomarker. These findings emphasize the potential of CYLD cleavage as a therapeutic target and a diagnostic marker of endotoxic shock.

## Results

### The cleavage of CYLD is required for LPS-induced endotoxic shock

Caspases are cysteine proteases that cleave specific substrates and play a pivotal role in programmed cell death and inflammation^16,17^. A previous study has demonstrated that Caspase8 cleaves the substrate CYLD at Asp215 to inhibit necroptosis in response to TNF^19^. Moreover, *in vitro* studies indicated that TLR4 activates Caspase8 to cleave CYLD in macrophages^20^. Notably, numerous studies have revealed that *Caspase8^-/-^Mlkl ^-/-^* or *Caspase8^-/-^Ripk3 ^-/-^* (or its adaptor, FADD-deficient) mice exhibit resistance to LPS-induced septic shock^12–15^. However, whether CYLD cleavage is involved in LPS-induced endotoxic shock remains unclear.

We first confirmed that LPS stimulation induces Caspase8-mediated cleavage of CYLD at D215, resulting in the formation of the N-terminal P25 fragment (CP25) in BMDMs **(Fig. S1a-d)**. To determine the *in vivo* role of CYLD cleavage in LPS-induced endotoxic shock, we generated knock-in mice with a point mutation at the 215 position by substituting aspartic acid with alanine (*Cyld^D^*^215^*^A/D^*^215^*^A^*mutant mice) **(Fig. S1e)**. *Cyld^D^*^215^*^A/D^*^215^*^A^* mice were viable and displayed normal development^30^, with CYLD protein levels in various tissues similar to those in wild-type mice **(Fig. S1f)**. Surprisingly, we found that *Cyld^D^*^215^*^A/D^*^215^*^A^* mice exhibited significant resistance to LPS-induced endotoxic shock compared to WT mice, with a higher survival rate **(Fig. 1a)**, reduced serum cytokine production (including IL-6,

**Figure 1.**
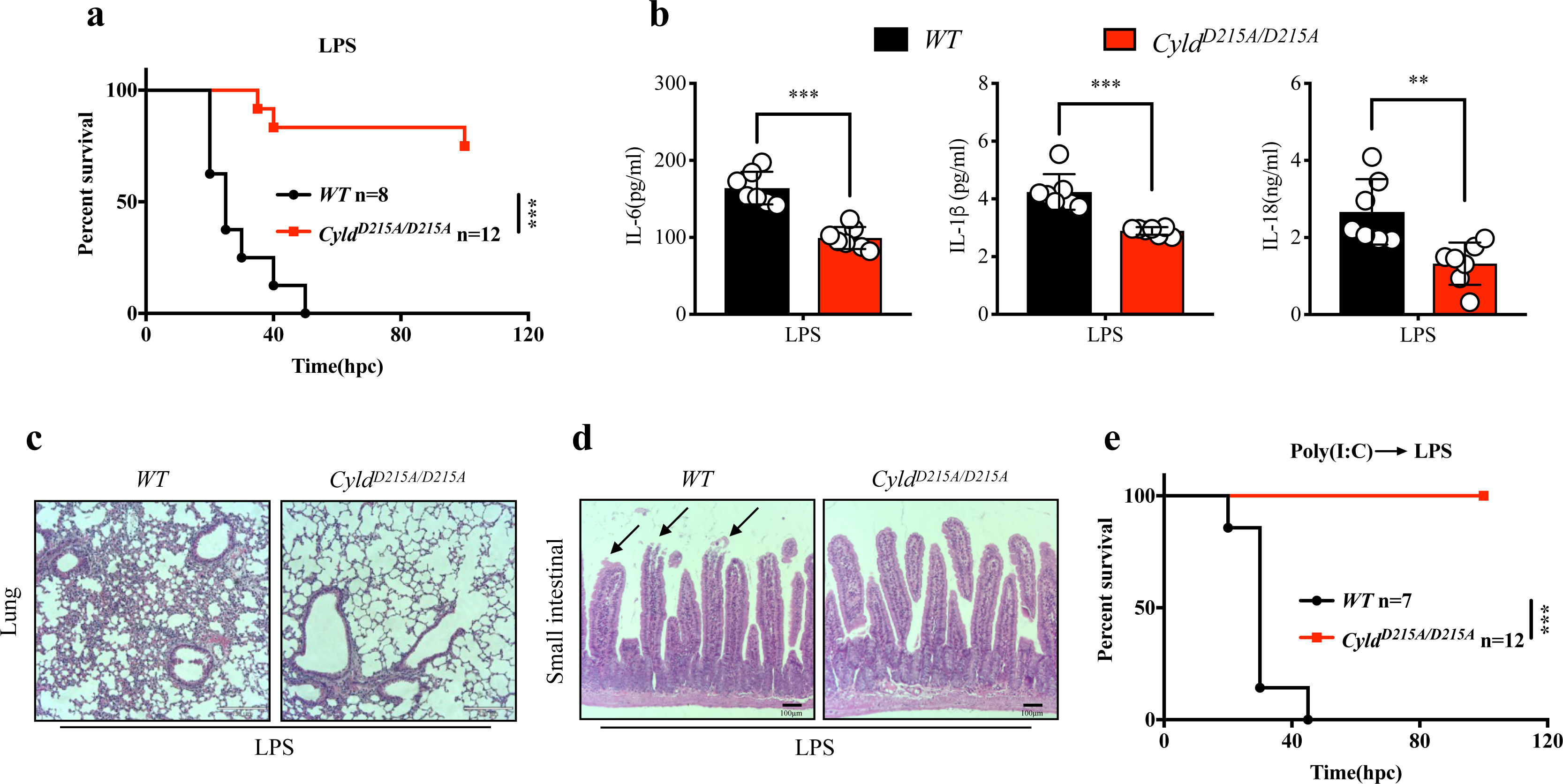
The cleavage of CYLD is required for LPS-induced endotoxic shock. **a**, Survival of WT (n=8) and *Cyld^D^*^215^*^A/D^*^215^*^A^* (n=12) mice challenged intraperitoneally with 50mg/kg LPS (O55:B5). **b**, Mice of indicated genotypes were injected intraperitoneally with 40mg/kg LPS (O55:B5). Serum was collected 12 hours after LPS treatment and subjected to ELISA analysis of IL-6, IL-1β and IL-18. **c-d,** Mice of indicated genotypes were injected intraperitoneally with 25mg/kg LPS (O55:B5). Lungs and small intestines were collected 9 hours after LPS treatment. The representative images of hematoxylin and eosin (H&E) stained sections are shown. **e,** Survival of WT (n=7) and *Cyld^D^*^215^*^A/D^*^215^*^A^* (n=12) mice primed with Poly(I:C) (LMW, 4 mg/kg) and then challenged intraperitoneally 6 hours later with LPS (O111:B4, 5 mg/kg). The P value of survival curve was determined by a log-rank (Mantel-Cox) test. P values calculated using t-test **(b)**. ^□^P<0.05, ^□□^P<0.01, ^□□□^P<0.001.

IL-1β, and IL-18), and milder tissue injury **(Fig. 1b-d)**. Consistently, we observed CYLD cleavage in various tissues, including the lungs, bone marrow, spleen, and liver, following LPS injection **(Fig. S1g)**. This observation was supported by a time-dependent reduction in full-length CYLD protein and a corresponding increase of CP25 **(Fig. S1g)**.

In addition to detecting of *Cyld^D^*^215^*^A/D^*^215^*^A^*mice resistant to LPS-induced endotoxic shock, we investigated the involvement of non-cleavable CYLD in TLR4-independent endotoxemia, a condition in where the Caspase11-mediated noncanonical inflammasome responds directly to LPS ^31^. We subjected both WT and *Cyld^D^*^215^*^A/D^*^215^*^A^*mice to priming with the TLR3 agonist Poly(I:C), followed by challenge with a low dose of LPS. The results showed that while none of the WT mice survived the challenge, most *Cyld^D^*^215^*^A/D^*^215^*^A^*mice survived **(Fig. 1e)**. These findings demonstrated that the cleavage of CYLD, mediated by Caspase8, plays a pivotal role in both canonical and noncanonical inflammasome-mediated endotoxic shock.

### CYLD plays a protective role downstream of Caspase8 in endotoxic shock

Although we found that *Cyld^D^*^215^*^A/D^*^215^*^A^* mice exhibited resistance to LPS-induced endotoxic shock, previous studies have shown that *Cyld* knock-out mice were more susceptible to LPS-induced endotoxic shock than WT mice. This paradox prompted us to speculate that the cleavage of CYLD by Caspase8 at position 215 facilitates CYLD degradation, subsequently promoting endotoxemia. To test this hypothesis, we confirmed the susceptibility of *Cyld^-/-^* mice to endotoxic shock at 20 hours post-LPS injection^32^, despite no significant difference in the survival curves eventually between WT and *Cyld^-/-^*mice **(Fig. 2a)**. Moreover, we observed no significant difference between WT and *Cyld^-/-^* mice in noncanonical septic shock induced by Poly(I:C) combined with LPS **(Fig. 2b)**. In addition, we investigated whether the resistance to endotoxic shock observed in *Caspase8^-/-^Mlkl^-/-^* mice was due to the absence of CYLD protein degradation mediated by Caspase8 cleavage at the D215 site. To address this, we generated *Caspase8^-/-^Mlkl^-/-^Cyld^-/-^* mice and challenged them with LPS or Poly(I:C) combined with LPS. Strikingly, the deletion of *Cyld* in *Caspase8^-/-^Mlkl^-/-^*mice led to a significant restoration of their susceptibility to endotoxic shock compared to *Caspase8^-/-^Mlkl^-/-^* mice **(Fig. 2c-d)**. These findings suggest that the activation-induced cleavage of CYLD by Caspase8 lead to the degradation and functional impairment of CYLD, thereby promoting LPS-induced endotoxic shock.

**Figure 2.**
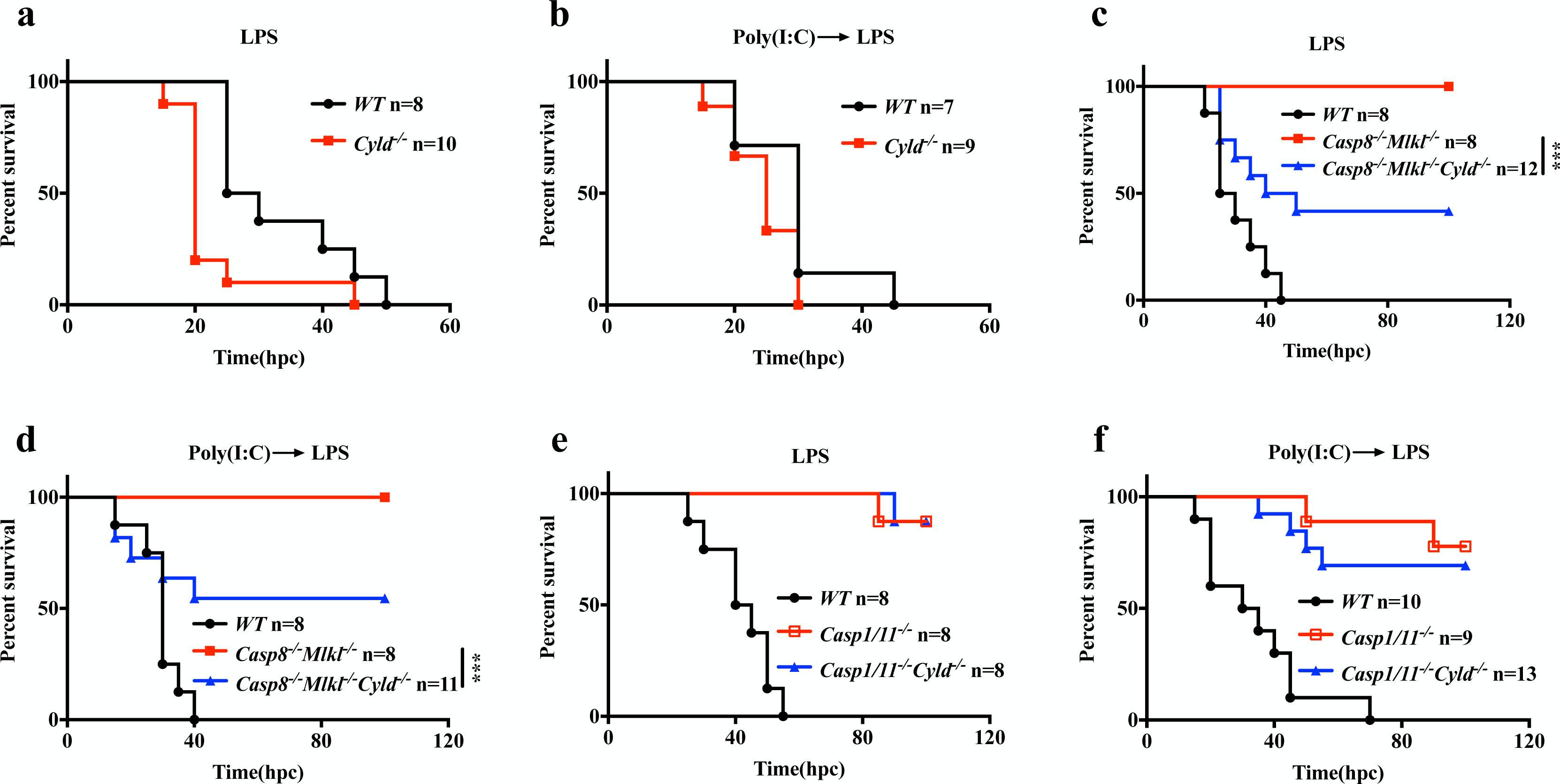
CYLD serves as an immunosuppressor downstream of Caspase8. **a**, Survival of WT (n=8) and *Cyld^-/-^* (n=10) mice challenged intraperitoneally with 50mg/kg LPS (O55:B5). **b**, Survival of WT (n=7) and *Cyld^-/-^* (n=9) mice primed with Poly(I:C) (LMW, 4 mg/kg) and then challenged intraperitoneally 6 hours later with LPS (O111:B4, 5 mg/kg). **c**, Survival of WT (n=8), *Casp8^-/-^Mlkl^-/-^* (n=8) and *Casp8^-/-^Mlkl^-/-^Cyld^-/-^* (n=12) mice challenged intraperitoneally with 50 mg/kg LPS (O55:B5). **d**, Survival of WT (n=8), *Casp8^-/-^Mlkl^-/-^*(n=8) and *Casp8^-/-^Mlkl^-/-^Cyld^-/-^* (n=11) mice primed with Poly(I:C) (LMW, 4 mg/kg) and then challenged intraperitoneally 6 hours later with LPS (O111:B4, 5 mg/kg). **e**, Survival of WT (n=8), *Casp1/11^-/-^* (n=8) and *Casp1/11^-/-^Cyld^-/-^*(n=8) mice challenged intraperitoneally with 50 mg/kg LPS (O55:B5). **f**, Survival of WT (n=10), *Casp1/11^-/-^* (n=9) and *Casp1/11^-/-^Cyld^-/-^* (n=13) mice primed with Poly(I:C) (LMW, 4 mg/kg) and then challenged intraperitoneally 6 hours later with LPS (O111:B4, 5 mg/kg). The P value of survival curve was determined by a log-rank (Mantel-Cox) test. ^D^P<0.05, ^DD^P<0.01, ^DDD^P<0.001.

Activation of Caspase1 and 11 in canonical and non-canonical inflammasomes, respectively, can contribute to pyroptosis^4^. *Caspase1/11^-/-^* mice exhibit substantial resistance to lethal endotoxic shock induced by LPS or Poly(I:C) combined with LPS. To investigate whether CYLD is involved in the *Caspase1/11^-/-^* mice in resistance to LPS-induced endotoxic shock, we generated *Caspase1/11^-/-^Cyld^-/-^* mice and challenged them with LPS or Poly(I:C) combined with LPS. Remarkably, both *Caspase1/11^-/-^Cyld^-/-^*and *Caspase1/11^-/-^* mice displayed comparable resistance to endotoxic shock **(Fig. 2e-f)**. These results suggest that during endotoxic shock induced by LPS or Poly(I:C), CYLD may function upstream of Caspase1/11.

### CYLD inhibits inflammasome activation by repressing the expression of both ***Nlrp3* and *Caspase11***

Next, we investigated how CYLD regulates canonical and non-canonical inflammasomes activation via cleavage at D215. We isolated WT and *Cyld^D^*^215^*^A/D^*^215^*^A^*BMDMs and treated them with various stimulators. In response to canonical inflammasome stimulations, *Cyld^D^*^215^*^A/D^*^215^*^A^* BMDMs showed reduced secretion of cleaved Casp1 and mature IL-1β compared to WT BMDMs **(Fig. 3a-c)**. Consistently, *Cyld^D^*^215^*^A/D^*^215^*^A^* BMDMs exhibited decreased levels of cleaved GSDMD and reduced cell death in response to non-canonical inflammasome stimulation compared to WT BMDMs **(Fig. 3d)**. These results indicate that the cleavage of CYLD at D215 leads to its degradation and functional impairment, thereby promoting both canonical and non-canonical inflammasomes activation.

**Figure 3.**
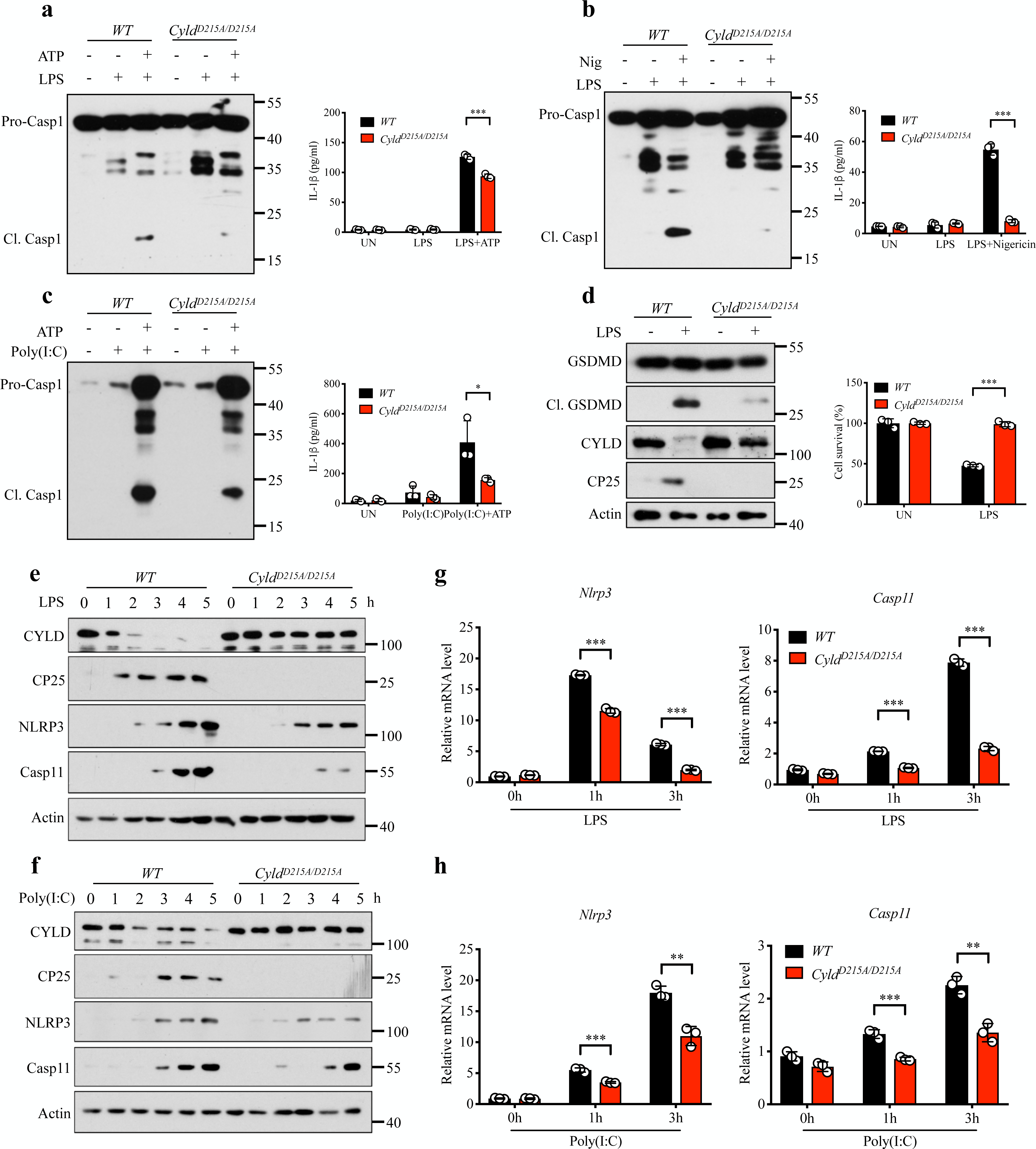
*Cyld^D^*^215^*^A/D^*^215^*^A^* inhibits the activation of inflammasome signaling pathways by negatively regulating the expression of *Caspase11* and *Nlrp3* genes. a-b, BMDMs were stimulated with LPS (100ng/ml, O55:B5) for 5 hours followed by ATP **(a)**, nigericin **(b)** for 30 min. culture supernatants were precipitated, and cell lysates were collected and analyzed for cleavage of procaspase1 and IL-1β. **c**, BMDMs were stimulated with Poly(I:C) (100μg/ml) for 5 hours followed by ATP for 30 min. culture supernatants were precipitated, and cell lysates were collected and analyzed for cleavage of procaspase1 and IL-1β. **d**, BMDMs were pretreated with LPS (100ng/ml, O55:B5) for 5 hours followed by LPS electroporation. Cell viability was determined using the Cell Titer-Glo kit. Cell lysates were collected and analyzed for cleavage of procaspase1 and IL-1β. **e-f**, Western blot of WT and *Cyld^D^*^215^*^A/D^*^215^*^A^* BMDMs stimulated with LPS (100ng/ml, O55:B5) **(e)** or Poly(I:C) (100μg/ml) **(f)** for indicated times. **g-h**, qPCR analysis of WT and *Cyld^D^*^215^*^A/D^*^215^*^A^* BMDMs stimulated with LPS (100ng/ml, O55:B5) **(g)** or Poly(I:C) (100μg/ml) **(h)** for indicated times. P values calculated using t-test **(a-d, g-h)**. ^D^P<0.05, ^DD^P<0.01, ^DDD^P<0.001.

Given that both canonical and non-canonical inflammasomes require a priming phase before activation^33^, we investigated whether CYLD cleavage plays a role in this priming phase. To explore this hypothesis, we treated WT and *Cyld^D^*^215^*^A/D^*^215^*^A^*BMDMs with LPS and Poly(I:C). Interestingly, we observed a significant reduction in the protein level of both NLRP3 and Caspase11, key components of inflammasome assembly, in *Cyld^D^*^215^*^A/D^*^215^*^A^* BMDMs **(Fig. 3e-f)**. This effect was further attributed to differences at the transcriptional level **(Fig. 3g-h)**. These findings suggest that non-cleavable CYLD suppresses the transcription of the key inflammasome components, *Nlrp3* and *Caspase11*, during the priming phase, thereby inhibiting inflammasome activation.

### CYLD inhibits NF-κB activation and inflammatory cytokines release

It has been reported that the transcription of *Nlrp3* and *Caspase11* is regulated by the NF-κB signaling pathway^34,35^. Considering the dampened transcription of *Nlrp3* and *Caspase11* in *Cyld^D^*^215^*^A/D^*^215^*^A^* BMDMs, we hypothesized that CYLD regulates *Nlrp3* and *Caspase11* transcription via the NF-κB pathway. To investigate this hypothesis, we assessed NF-κB signaling activation in *Cyld^D^*^215^*^A/D^*^215^*^A^* peritoneal macrophages (PMs) following LPS treatment. We observed a significant reduction in phosphorylated P65 in *Cyld^D^*^215^*^A/D^*^215^*^A^* PMs, with no significant impact on other components of the NF-κB pathway **(Fig. 4a)**. These findings correlated with the decreased production of TNFα and IL-6 in LPS-stimulated *Cyld^D^*^215^*^A/D^*^215^*^A^* PMs compared to that in WT PMs **(Fig. 4b-c)**. Given that the CYLD D215A mutant disrupted LPS-triggered TRIF-dependent CYLD cleavage, thus inhibiting CYLD degradation, we speculate that sustained CYLD protein levels are the primary reason for the attenuated NF-κB signaling in *Cyld^D^*^215^*^A/D^*^215^*^A^* PMs, because CYLD remains unprocessed and is not subject to degradation.

**Figure 4.**
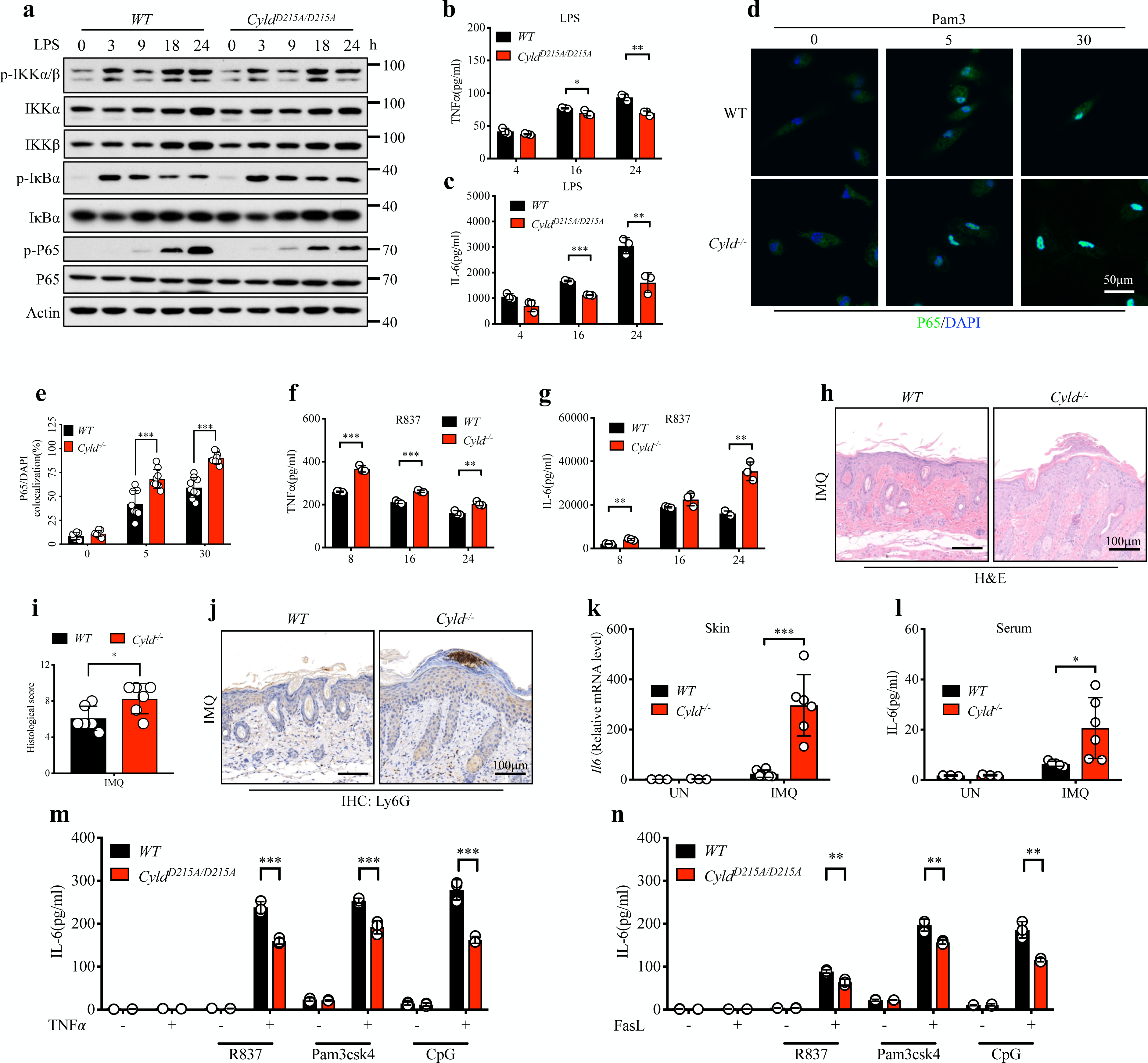
CYLD inhibits NF-κB activation and inflammatory cytokine release. **a**, Western blot of PM cells stimulated with LPS (100ng/ml, O55:B5) for indicated times. **b-c**, Plots show TNFα **(b)** and IL-6 **(c)** secretion of WT and *Cyld^D^*^215^*^A/D^*^215^*^A^* PMs treated with LPS (100ng/ml, O55:B5) for indicated times. **d-e**, Representative fluorescence micrographs (**d**) and quantification of the ratio of P65/DAPI colocalization (**e**) in WT and *Cyld^-/-^*PMs stimulated with Pmascsk4 (1ug/ml) for 0, 5, 30 minutes. Scale bars represent 50μm. P65 (green) (Cell Signaling Technology); Nucleus (blue). **f-g**, Plots show TNFα (**f**) and IL-6 (**g**) secretion of WT and *Cyld^-/-^* PMs treated with R837 (2μg/ml) for indicated times. **h,** Representative skin histological images of indicated mouse genotypes following daily topical treatment with imiquimod. Scale bars represent 100μm. **i**, Total histologic skin scores for indicated mouse genotypes following daily topical treatment with imiquimod. **j**, Representative Ly6G immunohistochemistry of skin for indicated mouse genotypes following daily topical treatment with imiquimod. Scale bars represent 100μm. **k**, RNAs from skins in i were isolated and used for expression analysis of IL6 using qPCR. **l**, Serum from indicated mouse in e was collected and subjected to ELISA analysis of IL-6. **m-n**, Plots show IL-6 secretion among BMDMs pretreated with TNFα (**m**) and FasL (**n**) for 24 hours before stimulated with the indicated TLR agonists for an additional 24 hours. P values calculated using t-test **(b-c, e-g, I, k-n)**. ^D^P<0.05, ^DD^P<0.01, ^DDD^P<0.001.

Because CYLD cleavage is TRIF-dependent, TRIF-independent stimuli, such as Pam3csk4 (TLR1/2 ligand), R837 (TLR7 ligand), and CpG (TLR9 ligand), do not induce CYLD cleavage **(Fig. S2a-b)**. If LPS or Poly(I:C) stimulation promotes NF-κB activation by cleaving and degrading CYLD, TRIF-independent stimuli should not decrease CYLD protein levels. As a result, this would lead to increased NF-κB activation in *Cyld^-/-^* cells compared to WT cells. To validate this, we assessed the responses to three TRIF-independent stimuli: Pam3csk4, R837, and CpG. As expected, Pam3csk4-treated *Cyld^-/-^* PMs displayed increased phosphorylation and nuclear translocation of P65 compared to WT cells **(Fig. 4d-e, S2c-d)**. Similarly, Pam3csk4 and R837 stimulation induced *Cyld^-/-^* PMs to produce higher levels of TNFα and IL-6 compared to WT PMs **(Fig. 4f-g, S2e-f)**. To extend these findings *in vivo*, we induced psoriasis in WT and *Cyld^-/-^* mice using imiquimod (R837) to trigger a

TRIF-independent inflammatory disease model. Histological analysis revealed exacerbated psoriatic lesions in *Cyld^-/-^* mice, characterized by epidermal hyperplasia and inflammatory cell infiltration **(Fig 4h-j)**, along with elevated skin expression of *IL-6* and serum IL-6 levels compared to that in WT mice **(Fig 3k-l)**. Collectively, these results demonstrate that TRIF-dependent stimuli activate Caspase8 to cleave CYLD, leading to its degradation and functional impairment, which promotes P65 phosphorylation and NF-κB activation, whereas TRIF-independent stimuli do not trigger CYLD cleavage.

Given that TNFα triggers Caspase8-dependent cleavage of CYLD in addition to TRIF-dependent manner, we speculated whether TNFα could potentially enhance NF-κB activation-mediated cytokine production downstream of TRIF-independent TLRs. Notably, we observed a minimal baseline presence of CP25 in WT BMDMs after LPS stimulation **(Fig. S2a-b, g)**, whereas CP25 was entirely absent in *Tnfr1^-/-^* BMDMs **(Fig. S2g)**, indicating a reliance of CYLD cleavage on cell-autocrine TNFα. Next, we verified that TNFα and FasL can also induce TRIF-independent, FADD/Caspase8-dependent CYLD cleavage **(Fig. S2h-k)**. To explore the involvement of CYLD cleavage in NF-κB signaling activation, we evaluated responses to TRIF-independent ligands (Pam3csk4, R837, and CpG) that do not cleave CYLD. These stimuli produced comparable IL-6 levels in the WT and *Cyld^D^*^215^*^A/D^*^215^*^A^* PMs. However, we found that pre-treatment with TNFα or FasL resulted in CYLD cleavage in WT PMs, amplifying IL-6 production in the presence of these TRIF-independent ligands more significantly in WT PMs than in *Cyld^D^*^215^*^A/D^*^215^*^A^* PMs **(Fig. 4m-n)**. Thus, these findings further emphasize the inhibitory function of the CYLD D215A mutant on the cytokine response by preserving a stable, full-length CYLD protein.

### CYLD regulates P65 activation through direct interaction with P65

The above analyses suggest that CYLD deletion or the D215A mutation affects NF-kB activation by modulating P65 phosphorylation. Next, we investigated whether CYLD directly interact with P65. Notably, CYLD efficiently co-immunoprecipitated with P65 in transfected 293T cells **(Fig. 5a-b)**. This interaction was further supported by immunofluorescence assays, which confirmed their co-localization **(Fig. 5c)**. Moreover, through molecular mapping of truncated forms of CYLD and P65 mutants, we identified the CYLD CG3 and DUB domains (amino acids 472–540 and 584-955) as the region responsible for binding to amino acids 194–255 of P65 **(Fig. 5d-g)**.

**Figure 5.**
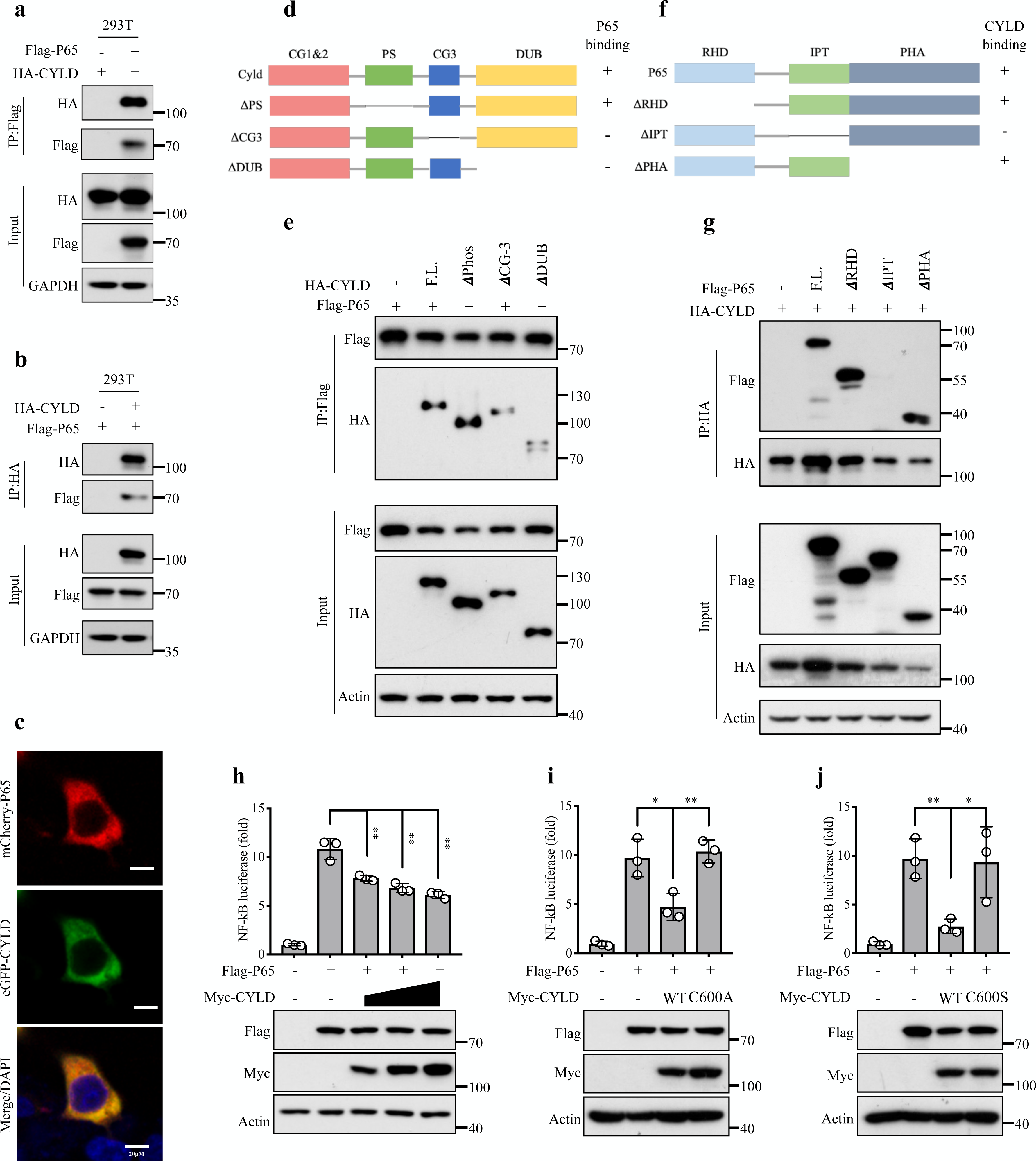
CYLD regulates P65 activation through direct interaction with P65. **a-b**, 293T cells were transfected with plasmids encoding HA-CYLD, Flag-P65 and corresponding empty vector, followed by immunoprecipitation with anti-Flag beads **(a)** and anti-HA beads **(b)** and followed by immunoblotting analysis. **c**, 293T cells were transfected with plasmids encoding GFP-CYLD and mCherry-P65, and co-localization of P65 and CYLD was detected by immunofluorescence. **d**, Schematic of the full-length CYLD and its deletion mutations. **e**, Cell lysates derived from 293T cells transfected with Flag-P65 and empty vector, HA-tagged CYLD or CYLD truncation mutants were analyzed by co-immunoprecipitation (co-IP) assay with Flag-beads, followed by immunoblotting. **f**, Schematic of the full-length P65 and its deletion mutations. **g**, Cell lysates derived from 293T cells transfected with HA-CYLD and empty vector, Flag-tagged P65 or P65 truncation mutants were analyzed by co-immunoprecipitation (co-IP) assay with Flag-beads, followed by immunoblotting. **h**, 293T cells were transfected with NF-κB luciferase reporter together with Flag-P65, Myc-CYLD (with increased dosage), empty vector and pRL-TK. NF-κB activity was assayed. **i-j**, 293T cells were transfected with NF-κB luciferase reporter together with empty vector and pRL-TK, Flag-P65, Myc-CYLD and its mutants Myc-CYLD-C600A **(i)** and Myc-CYLD-C600S **(j)**. Then NF-κB activity was assayed. P values calculated using t-test **(h-j)**. ^D^P<0.05, ^DD^P<0.01, ^DDD^P<0.001.

Given that CYLD is a deubiquitinating enzyme, we examined whether the regulation of P65 activation relies on its enzymatic activity. By employing the NF-κB reporter assay system, we observed a dose-dependent suppressive effect of CYLD on P65 activation **(Fig 5h)**. However, this inhibitory effect was diminished upon mutation of the key amino acid (C600A or S), which is critical for CYLD enzymatic activity **(Fig 5i-j)**. Collectively, these findings demonstrate that CYLD regulates P65 activation through direct interaction with P65, and this regulatory function is dependent on the enzymatic activity of CYLD.

### CYLD, functioning as a deubiquitinase, specifically removes M1-linked ubiquitination from P65 to suppress its activation

We further explored the underlying mechanisms by which CYLD regulates P65 phosphorylation via its interaction with P65. Phosphorylation of P65 in LPS-stimulated J774 cells was accompanied by ubiquitination, a phenomenon also observed in PMs stimulated with Pam3csk4 **(Fig. S3a-b)**. This parallel occurrence indicates a significant correlation between P65 activation and ubiquitination status. Given the established role of CYLD as a deubiquitinase, we determined whether P65 serves as a functional substrate for CYLD-mediated deubiquitylation. Through co-expression experiments involving CYLD and P65 in 293T cells, we found that CYLD significantly reduced ubiquitination of P65 **(Fig. 6a)**. To investigate the specific ubiquitination types of P65 regulated by CYLD, we performed co-expression experiments with CYLD, P65, and distinct ubiquitination types, including M1, K6, K11, K27, K29, K33, K48, and K63-linked ubiquitination, in 293T cells. CYLD specifically removes M1-linked ubiquitination of P65, whereas it has no effect on other forms of ubiquitination **(Fig 6a, S3c-d)**. Consistent with this result, CYLD can reduce K48R- and K63R-linked ubiquitination of P65 but failed to remove M1A (non-M1)-linked polyubiquitin chains **(Fig. 6b, S3c)**. Moreover, catalytically inactive CYLD mutants (C600A or C600S) were unable to effectively reduce the total and M1-linked ubiquitination of P65 **(Fig. 6c-d, S3e-f)**. In contrast, the CRISPR/Cas9-mediated knockout of *Cyld* resulted in increased total and M1-linked ubiquitination of flag-P65 in both Hela and 293T cells **(Fig. 6e-f, S3g-h)**. Consistently, LPS-stimulated *Cyld^D^*^215^*^A/D^*^215^*^A^* PMs displayed diminished P65 ubiquitination in comparison to WT PMs, consistent with the noted decrease in P65 phosphorylation **(Fig. 6g)**. In addition, increased levels of ubiquitinated and activated P65 were observed in *Cyld^-/-^*PMs compared to those in WT PMs following Pam3csk4 stimulation **(Fig. 6h)**. Collectively, these findings demonstrate that CYLD functions as a specific DUB for the M1-linked ubiquitination of P65, thereby modulating its activation.

**Figure 6.**
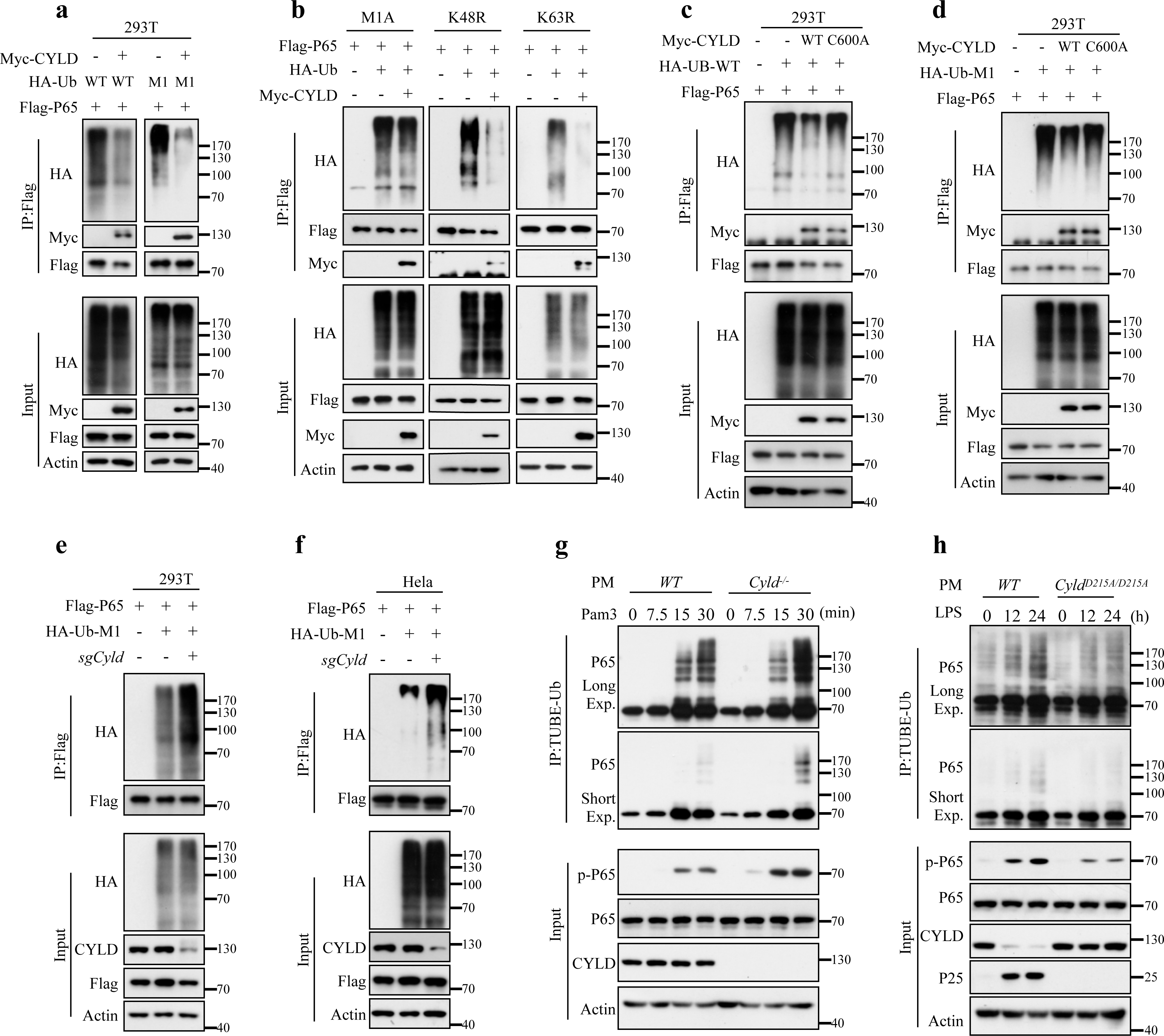
CYLD, functioning as a deubiquitinase, specifically removes M1-linked ubiquitination from P65 to modulate P65 activation. **a**, 293T cells were transfected with Flag-P65, Myc-CYLD, HA-WT and HA-M1 plasmids with the indicated combinations for 24h. The cell lysates were subjected to immunoprecipitation with anti-Flag beads and immunoblotted with the indicated antibodies. **b**, 293T cells were transfected with Flag-P65, Myc-CYLD, HA-M1A, HA-K48R and HA-K63R plasmids with the indicated combinations for 24h. The cell lysates were subjected to immunoprecipitation with anti-Flag beads and immunoblotted with the indicated antibodies. **c-d**, 293T cells were transfected with Flag-P65, Myc-CYLD, Myc-CYLD-C600A, HA-WT **(c)** and HA-M1 **(d)** plasmids with the indicated combinations for 24h. The cell lysates were subjected to immunoprecipitation with anti-Flag beads and immunoblotted with the indicated antibodies. **e-f**, 293T **(e)** and Hela **(f)** cells stably expressing control or *Cyld* small-guide RNA. And then cells were transfected with Flag-P65 and HA-M1 plasmids as indicated for 24h. The cell lysates were subjected to immunoprecipitation with anti-Flag beads and immunoblotted with the indicated antibodies. **g-h**, WT and *Cyld^-/-^* PMs were stimulated with Pmascsk4 (1μg/ml) for indicated times **(g)** and WT and *Cyld^D^*^215^*^A/D^*^215^*^A^* PMs treated with LPS (100ng/ml, O55:B5) for indicated times **(h)**. Cell lysates were subjected to immunoprecipitation using TUBE antibody and Flag-beads. TUBE-Ub samples and cell lysates (input) were analyzed using western blot.

### CYLD, in coordination with the LUBAC complex, modulates P65 M1-linked ubiquitination at K301/K303

The balance between ubiquitylation and deubiquitylation is tightly regulated by ubiquitin ligases and deubiquitinases^36^. Our results demonstrated that CYLD functions as a specific deubiquitinase (DUB) for the M1-linked ubiquitination of P65. In addition, the Lubac complex, composed of HOIP, HOIL-1, and SHARPIN subunits, has been identified as the only E3 ubiquitin ligase that catalyzes the formation of M1-linked ubiquitin chains^22^. Thus, we investigated whether the Lubac complex is involved in the ubiquitination of P65. To address this, we conducted co-expression experiments involving components of the Lubac complex and P65 in 293T cells, revealing direct interactions between all three Lubac subunits and P65 **(Fig. S4a)**. Moreover, the Lubac linear ubiquitination chain assembly complex exhibited a marked increase in the total and M1-linked ubiquitination of flag-P65 **(Fig. 7a)**. Intriguingly, the mutation of ubiquitin E3 ligase-inactive mutants of HOIP (C879A and C910A) weakened the enhancement of the ubiquitination of P65 **(Fig. 7b)**. Collectively, these findings demonstrated the regulatory role of the Lubac complex as an E3 ubiquitin ligase that directly interacts with P65 to modulate its ubiquitination status.

**Figure 7.**
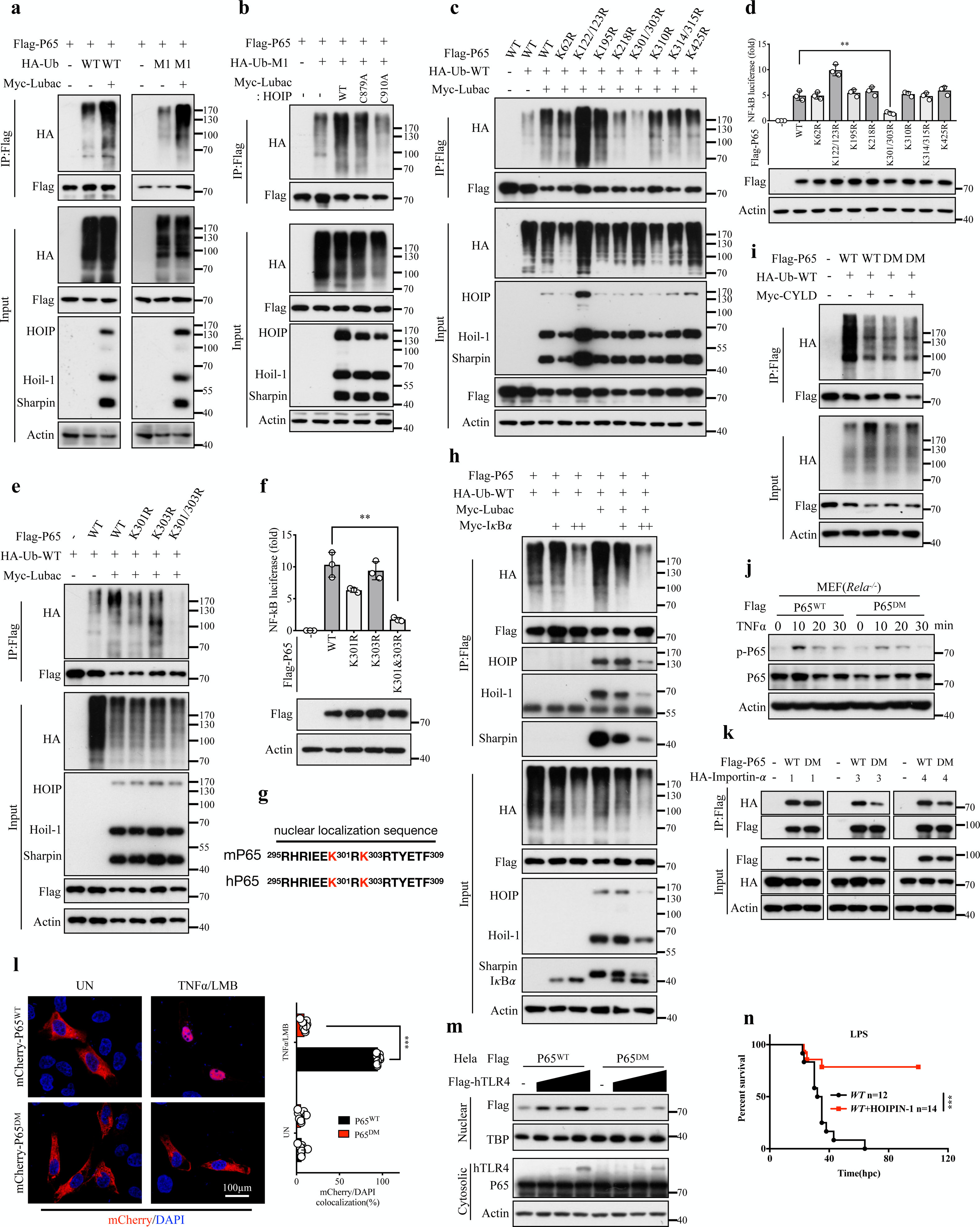
CYLD, in coordination with the LUBAC complex, modulates P65 nuclear translocation by regulating the M1-linked ubiquitin modification. **a**, 293T cells were transfected with Flag-P65, Myc-Lubac (including Myc-HOIP, Myc-HOIL-1 and Myc-Sharpin), HA-WT and HA-M1 plasmids with the indicated combinations. The cell lysates were subjected to immunoprecipitation with anti-Flag beads and immunoblotted with the indicated antibodies. **b**, 293T cells were transfected with Flag-P65, HA-M1, and Myc-Lubac (including Myc-HOIL-1 and Myc-Sharpin, Myc-HOIP and HOIP mutants Myc-HOIP-C879A, Myc-HOIP-C910A), and plasmids with the indicated combinations. The cell lysates were subjected to immunoprecipitation with anti-Flag beads and immunoblotted with the indicated antibodies. **c**, Hela cells were transfected with HA-WT, Myc-Lubac, Flag-P65 (and its mutants) and plasmids with the indicated combinations. The cell lysates were subjected to immunoprecipitation with anti-Flag beads and immunoblotted with the indicated antibodies. **d**, Hela cells were transfected with pRL-TK, NF-κB luciferase reporter, empty vector, Flag-P65 and its mutans. Then NF-κB activity was assayed. **e**, Hela cells were transfected with HA-WT, Myc-Lubac, Flag-P65 (and its mutants) and plasmids with the indicated combinations. The cell lysates were subjected to immunoprecipitation with anti-Flag beads and immunoblotted with the indicated antibodies. **f**, Hela cells were transfected with pRL-TK, NF-κB luciferase reporter, empty vector, Flag-P65 and its mutants. Then NF-κB activity was assayed. **g**, The nuclear localization sequences (NLS) in mP65 and hP65. With conserved lysine residues located in NLS indicated in red. **h**, Hela cells were transfected with HA-WT, Myc-Lubac, Myc-IκBα (with increased dosage), Flag-P65 and plasmids with the indicated combinations. The cell lysates were subjected to immunoprecipitation with anti-Flag beads and immunoblotted with the indicated antibodies. **i**, Hela cells were transfected with HA-WT, Myc-CYLD, Flag-P65 and the mutants and plasmids with the indicated combinations. The cell lysates were subjected to immunoprecipitation with anti-Flag beads and immunoblotted with the indicated antibodies. **j**, Immortalized *Rela^-/-^* MEF cells were transfected with Flag-P65 and its mutant Flag-P65-DM for 24h. Then cells were stimulated with mTNFα for indicated times and the cell lysates were analyzed using western blotting. **k**, Hela cells were transfected with HA-importin α1, HA-importin α3, HA-importin α4, Flag-P65 and the mutant Flag-P65^DM^ and plasmids with the indicated combinations. The cell lysates were subjected to immunoprecipitation with anti-Flag beads and immunoblotted with the indicated antibodies. **l**, Hela cells were transfected with plasmids encoding mCherry-P65 and mCherry-P65^DM^ for 24h. Then cells were pretreated with LMB (2ng/ml) for 20min followed by stimulated with mTNFα for 25min. The P65 was detected by immunofluorescence and the entry into the nuclear of P65 are quantified. **m**, Hela cells were transfected with Flag-hTLR4 (with increased dosage), Flag-P65 and Flag-P65^DM^ with the indicated combinations for 24h. Then stimulated the cells with LPS (100ng/ml) for 15min and separate the nucleus and cytoplasm by Nuclear and Cytoplasmic Protein Extraction Kit and perform subsequent Western blot. **n**, Survival of WT mice primed with saline or HOIPIN-1 (3mg/kg) and then challenged 6 hours later with LPS (O55:B5, 50mg/kg) intraperitoneally. The P value of survival curve was determined by a log-rank (Mantel-Cox) test. P values calculated using t-test **(d, f and l)**. ^D^P<0.05, ^DD^P<0.01, ^DDD^P<0.001.

To further elucidate the mechanism by which CYLD and the Lubac complex synergistically regulate the ubiquitination balance of P65, we identified the potential ubiquitination sites on P65 modified by CYLD and the Lubac complex. Using group-based prediction system for ubiquitin E3 ligase-substrate relations (GPS-Uber) (http://bdmpub.biocuckoo.org/prediction.php), we predicted 11 possible ubiquitination sites on P65 **(Fig. S4b)**. Subsequently, we systematically introduced mutations at each of these predicted sites by replacing lysine (K) with arginine (R) to evaluate their impact on P65 ubiquitination. Notably, we found that simultaneous mutations of K301 and K303 (referred to as P65^DM^) effectively diminished P65 ubiquitination **(Fig. S4c)**. Notably, alterations in K301 and K303 eliminated the intensified ubiquitination of flag-P65 catalyzed by the Lubac complex **(Fig. 7c)**. Consistently, we observed that mutations in K301 and K303 significantly suppressed NF-κB reporter activation in both 293T and Hela cells **(Fig. 7d, S4d)**.

To determine whether individual mutations at these two sites were sufficient to impact P65 ubiquitination, we introduced mutations of K301 and K303 to arginine (R) individually. However, our observations revealed that unlike the combined mutations at these two sites, which were critical in effectively suppressing both P65 ubiquitination and activation, single-site mutations in K301R or K303R did not significantly affect P65 ubiquitination and activation **(Fig. 7e-f, S4e)**. Next, we determined the timing of Lubac complex-mediated ubiquitination of P65. Since K301/K303 are located on the nuclear localization sequence (NLS) of P65 **(Fig. 7g)** and were shielded by IκBα. We speculated that the Lubac-mediated ubiquitination of P65 occurs after the dissociation of P65 from I_κ_Bα. Co-transfection experiments revealed that along with IκBα expression increased, more of nuclear localization sequence of P65 covered, P65 ubiquitination decreased **(Fig. 7h)**. This suggests that Lubac-mediated ubiquitination of P65 occurs after the dissociation of P65 from IκBα.

Furthermore, the dual mutation K301R/K303R (referred to as P65^DM^) was able to diminish the influence of CYLD on P65 linear ubiquitination, suggesting that CYLD specifically targets the removal of P65 linear ubiquitination at the specific K301 and K303 sites **(Fig. 7i)**. Consistent with these findings, the inhibitory effect of P65^DM^ on P65 activation was confirmed by a significant reduction in TNFα-triggered P65 phosphorylation **(Fig. 7j)**. As previously mentioned, K301/K303 are located within the NLS **(Fig. 7g)**, with importin α3 and α4 acting as the primary importin α isoforms facilitating P65 nucleus translocation^37,38^. Therefore, we next investigated whether P65^DM^ affects its interaction with these importin α proteins. We observed that the P65^DM^ mutation diminishes binding to importin α3 and α4, whereas retaining binding to importin α1 **(Fig. 7k)**. Accordingly, we observed a significant decrease in P65^DM^ nuclear translocation induced by TNFα or co-transfected with TLR4 before stimulated with LPS **(Fig. 7l-m, S4f)**, confirming that the M1-linked ubiquitination on K301/303 sites of P65 facilitates its nuclear translocation.

To further validate the critical role of P65 linear ubiquitination modulated by CYLD and the Lubac complex *in vivo*, we pre-treated WT mice with HOIPIN-1, a HOIP inhibitor, followed by LPS challenge to induce endotoxic shock. Remarkably, HOIPIN-1 pre-treatment resulted in a significant reduction in mortality in WT mice **(Fig. 7n)**. Collectively, these results demonstrated that CYLD, in collaboration with the LUBAC complex, specifically regulates M1-linked ubiquitin modification at the K301/303 residues of P65 downstream of detaching from IκBα, thereby modulating the nuclear translocation and activation of P65.

### CP25 in the serum could potentially serve as a biomarker for endotoxic shock and related inflammatory diseases

Although we established the requirement for CYLD cleavage at D215 by Caspase8 during endotoxic shock, the fate of the cleaved CP25 remained elusive. To explore whether CP25 can be secreted from the cytosol into the outside cells, we analyzed the presence of CP25 in both cell lysates and supernatants of bone marrow-derived macrophages (BMDMs) at different time points following LPS stimulation. Notably, we observed the presence of CP25 in the supernatant, and its levels gradually increased over time upon LPS stimulation. In contrast, *Cyld^D^*^215^*^A/D^*^215^*^A^*mutant cells, which did not produce CP25, exhibited only small amounts of full-length CYLD and completely lacked CP25 in the supernatant **(Fig. 8a)**. Thus, we also confirmed that CP25 does not undergo nuclear translocation but is secreted to the outside cells **(Fig. S5a)**.

**Figure 8.**
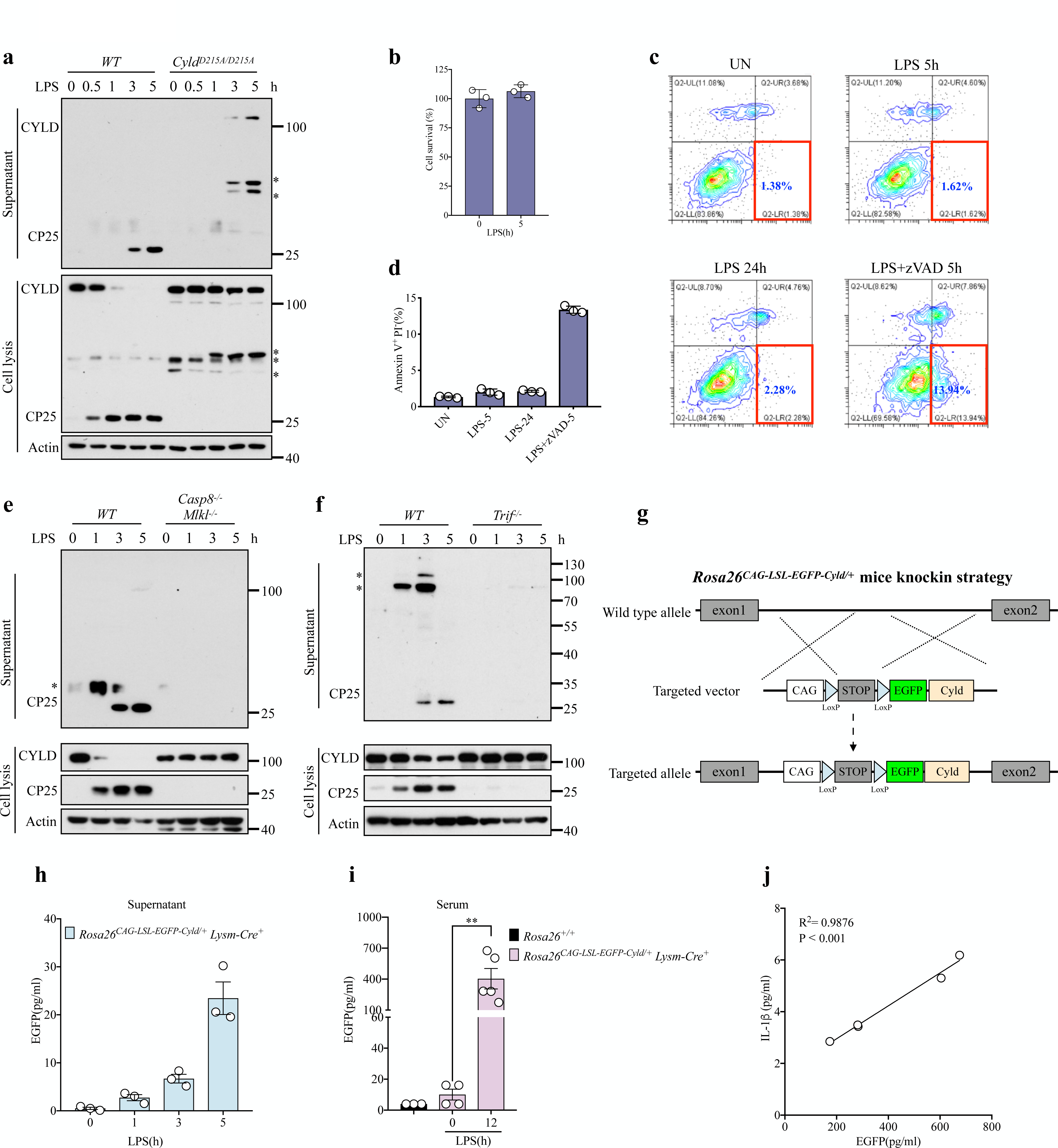
CP25 in the serum could potentially serve as a biomarker for endotoxic shock and related inflammatory diseases. **a**, WT and *Cyld^D^*^215^*^A/D^*^215^*^A^* BMDMs stimulated with LPS (100ng/ml, O55:B5) for indicated times. Cell lysates and supernatants were analyzed by Western blotting. **b**, BMDMs were treated with LPS (100ng/ml, O55:B5) for 5 hours. Cell viability was determined using the Cell Titer-Glo kit. **c-d**, Flow cytometric analysis (**c**) and quantifications (**d**) of BMDMs treated with LPS (100ng/ml, O55:B5) and LPS+zVAD for indicated times. Cells were stained with Annexin V and Propidium Iodide (PI) at the indicated time points. **e-f**, WT and *Casp8^-/-^Mlkl^-/-^* (**e**); WT and *Trif^-/-^* (**f**) BMDMs stimulated with LPS (100ng/ml, O55:B5) for indicated times. Cell lysates and supernatants were analyzed by Western blotting. **g**, Schematic showing the construction strategy of *Rosa26^CAG-LSL-EGFP-Cyld/+^*knock in mice. **h**, Plots show EGFP of *Lysm-Cre^+^ Rosa26^CAG-LSL-EGFP-Cyld/+^* BMDMs treated with LPS (100ng/ml, O55:B5) for indicated times. **i**, Plots show EGFP of WT and *Lysm-Cre^+^ Rosa26^CAG-LSL-EGFP-Cyld/+^*mice injected intraperitoneally with 25mg/kg of LPS (O55:B5). **j**, Correlations between serum EGPE-CP25 and IL-1β. P values calculated using t-test **(b, d, h and i)**. The P values was determined by linear regression **(i)**. ^D^P<0.05, ^DD^P<0.01, ^DDD^P<0.001. The asterisks represent the non-specific bands.

Next, we investigated the mechanisms underlying CP25 secretion. As CP25 lacks the signal peptide sequence required for protein secretion^39–41^, we initially investigated the involvement of the unconventional secretory pathway (the exosome-mediated^42^) in CP25 secretion. However, we found that the secretion of the CP25 is independent of this secretory pathway **(Fig. S5b)**. Moreover, we detected no cell death in WT BMDMs after LPS stimulation, indicating that CP25 secretion does not depend on membrane rupture caused by cell death **(Fig. 8b-d)**. Notably, neither CP25 nor noncleaved full-length CYLD was detected in the supernatant of *Caspase8^-/-^Mlkl^-/-^* BMDMs following LPS stimulation **(Fig. 8e)**. This observation suggests the essential role of Caspase8 in the secretion of both CP25 and full-length CYLD. Because the upstream protein TRIF is required for the LPS-stimulated cleavage activity of Caspase8, we assessed whether CYLD secretion was dependent on TRIF. We found that full-length CYLD in LPS-stimulated *Trif^-/-^* BMDMs remained in the cytosol and could not be secreted **(Fig. 8f)**. Thus, we propose that the secretion of CP25 depends on the recruitment of the TRIF/Caspase8 complex facilitated by the TLR4 receptor upon LPS stimulation, which establishes a novel secretory pathway dependent on the formation of a secretory complex within the cell cytosol.

Given that CP25 can be secreted extracellularly upon LPS stimulation, and that secretion increases with stimulation duration, we next investigated whether CP25 serves as a potential disease biomarker detectable in the serum. Owing to the lack of available ELISA reagents for quantifying serum CP25 levels, we generated a mouse line expressing an EGFP-CYLD fusion protein (*Rosa26^CAG-LSL-EGFP-Cyld/+^*) by introducing an EGFP tag at the N-terminus of CYLD behind the CAG-LSL elements between exon 1 and exon 2 **(Fig. S5c)**. Then we generated the *Lysm-Cre*^+^ Rosa26^CAG-LSL-EGFP-Cyld/+^ mice by crossing Rosa26^CAG-LSL-EGFP-Cyld/+^ mice with *Lysm-Cre^+^* mice, thereby inducing EGFP-CYLD with specific expression in monocytes/macrophages **(Fig. 8g)**. The *Lysm-Cre*^+^ *Rosa26^CAG-LSL-EGFP-Cyld/+^*mice were healthy and exhibited normal growth. To validate extracellular secretion of EGFP- CP25, BMDMs isolated from *Rosa26^CAG-LSL-EGFP-Cyld/+^Lysm-Cre^+^*mice were stimulated with LPS. Then, EGFP-CP25 was identified in the supernatant, and its amount increased progressively with time **(Fig. 8h)**. Next, we assessed whether EGFP-CP25 could be detected in serum during LPS-induced endotoxic shock. *Rosa26^CAG-LSL-EGFP-Cyld/+^Lysm-Cre^+^*mice were injected with LPS, and a significant elevation of EGFP-CP25 was observed in the serum of LPS-injected mice compared to non-injected mice using a GFP ELISA kit **(Fig. 8i)**. These findings suggest that the CP25 can be secreted from cells and detected in the serum of mice experiencing LPS-induced endotoxic shock. Moreover, the EGFP-CP25 level has strongly correlated with corresponding serum IL-1β **(Fig. 8j)**. This implies that the CP25 in serum may serve as a potential biomarker of endotoxic shock and related inflammatory diseases.

## Discussion

Caspase8 is a key regulator of apoptosis, necroptosis, pyroptosis and the inflammatory response^16,17^. Previous studies have demonstrated that *Caspase8^-/-^Mlkl^-^*^/-^(or *Caspase8^-/-^Ripk3^-/-^*) mice exhibit resistance against LPS or gram-negative bacteria-induced endotoxic shock, with this resistance primarily attributed to Caspase8 rather than the RIPK3/MLKL axis^12–15^. In this study, we performed genetic analysis delineating the role of Caspase8 mediated CYLD cleavage in LPS induced endotoxic shock. Our results define the requirement for Caspase8 mediated cleavage of CYLD in inflammasome activation and inflammatory cytokine production. We demonstrated that Caspase8 cleaves the substrate protein CYLD at residue D215, leading to the loss of intact CYLD, reduced deubiquitinating of P65 and increased NF-κB activation. Furthermore, NF-κB activation promotes the expression of *Caspase11* and *Nlrp3* during priming stages^12,33^, consequently leading to inflammasomes activation; on the other hand, the activation of NF-κB also promotes the production of inflammatory cytokines^43,44^, which exacerbate LPS induced endotoxic shock **(Fig. S5d)**. Consistent with previous studies that FADD and Caspase8 regulate the priming stages of both canonical and non-canonical inflammasomes^12,45^, thereby involving in the regulation of endotoxic shock, we found that Caspase8 plays a critical role in the priming stage by the cleaving and reducing CYLD, which was confirmed by *Cyld^D^*^215^*^A/D^*^215^*^A^* macrophages with weaker expression of *Nlrp3* and *Caspase11*. It has been shown previously that Caspase8 plays a pivotal role in inducing pro-inflammatory gene expression, depending on its enzymatic activity, under bacterial infection or TLR stimulation^13^. In addition, we demonstrated that Caspase8, through its enzymatic function, cleaves CYLD to regulate the inflammatory response. Accordingly, the knockout of the *Cyld* gene in *Caspase8^-/-^Mlkl^-/-^*mice significantly restored sensitivity to endotoxic shock, emphasizing the importance of maintaining stable CYLD levels in *Caspase8^-/-^Mlkl^-/-^* mice for resistance to endotoxic shock. Therefore, our study revealed that CYLD is a critical substrate of Caspase8, and cleavage at D215 plays a pivotal role in endotoxic shock.

It is worth noting that, despite knocking out *Cyld* in *Caspase8^-/-^Mlkl^-/-^* mice restored sensitivity to LPS-induced endotoxic shock, the sensitivity of *Caspase8^-/-^Mlkl^-/-^Cyld^-/-^* mice to endotoxic shock remained weak compared to that of WT mice. This may be caused by the following three reasons: firstly, pro-apoptotic function of Caspase8 within the lower small intestine contributes to this process^14^; secondly, Caspase8 has also been implicated in the cleavage of GSDMD and inflammasome regulation^46–48^; thirdly, activated Caspase8 can cleave various substrates to promote cytokine production in addition to CYLD, such as N4BP1 or other undefined substrates^13,18^. Further investigations, such as concurrent *Cyld* and *N4bp1* knockout in the *Caspase8^-/-^Mlkl^-/-^* background, might largely restore sensitivity to endotoxic shock.

CYLD has been established as a DUB that targets various NF-kB signaling factors, including NEMO^26–28^, TRFA2^26–28^, TRAF6^26–28,49^, TRAF7^49^, Bcl-3^29^, RIP1^50^, and TAK1^51^. However, the specific targets for deubiquitination vary depending on the stimulus conditions and the cell type^52^. The specific mechanism by which CYLD negatively regulates the NF-kB signaling pathway remains unclear and warrants further research. In this study, we found that under LPS stimulation, p-P65 in WT macrophages was stronger than that in *Cyld^D^*^215^*^A/D^*^215^*^A^*macrophages, whereas the phosphorylation and degradation of IκBα have no significant difference between the two genotypes. Moreover, under Pam3csk4 stimulation, which can not induce cleavage of CYLD, p-P65 in WT was weaker than in *Cyld^-/-^* macrophages, but the phosphorylation and degradation of IκBα were not significantly different between the two genotypes. This suggests that CYLD is involved in the regulation of P65. Furthermore, we found that CYLD inhibited the activation and entry into nuclear of P65 by directly interacting with and removing the M1 type ubiquitin chain connected to P65.

Ubiquitination plays a pivotal role in the regulation of P65, and previous studies focused on the degradation of P65 mediated by ubiquitination^53–56^. Our research unravels a novel mechanism involving M1-linked polyubiquitination for nuclear entry of P65 mediated NF-kB activation. We found that Lubac complex catalyzed M1 linkage on P65 after dissociation with IκBα. Subsequently, CYLD removed the linear ubiquitin chain from P65 and influenced its entry into the cytosol and phosphorylation. Additionally, we systematically screened potential ubiquitination modification sites and ultimately identified K301 and K303 on P65 as the specific targets for linear ubiquitination. The K301/303 sites are located within the nuclear localization sequence (NLS) of P65. Mutations changing K to R at these positions resulted in reduced ubiquitination, affecting binding of P65 to importin α3 and importin α4, thus reducing its nuclear translocation. Collectively, the identification of these critical deubiquitination sites not only enriches our understanding of the ubiquitination regulatory mechanisms governing P65-mediated NF-kB activation, but also provides novel targets for the development of specific drugs targeting P65 ubiquitination regulation.

In this study, we generated CYLD D215A mutant mice (*Cyld^D^*^215^*^A/D^*^215^*^A^*) resistant to Caspase8 cleavage. *Cyld^D^*^215^*^A/D^*^215^*^A^* mice demonstrated significant resistance to lethal endotoxic shock. Notably, the removal of *Cyld* in *Caspase8^-/-^Mlkl^-/-^* mice effectively restored their sensitivity to endotoxic shock, confirming that the maintenance of CYLD stability is the crucial factor responsible for the observed resistance in *Caspase8^-/-^Mlkl^-/-^* mice. Based on these findings, the development of drugs specifically targets the cleavage of CYLD at the Asp215 site holds promise for the prevention and treatment of endotoxemia. Furthermore, we unexpectedly found that CP25, the N-terminal cleavage fragment of CYLD, was secreted extracellularly via a pathway mediated by the TRIF/Caspase8 complex-a novel secretion mechanism. Notably, septic mice challenged with LPS exhibited increased serum CYLD-CP25 levels over time, indicating that this CP25 fragment may serve as a novel inflammatory disease marker that can be detected through blood analysis. Therefore, Caspase8-mediated cleavage of CYLD at D215 is not only a potentially critical target in inflammatory conditions such as endotoxic shock but the resulting CP25 fragment may also emerge as a new blood based inflammatory biomarker for these diseases.

## Methods

### Mice

The C57BL/6J mice were housed in a specific pathogen-free (SPF) facility. *Caspase8^+/-^*^57^, *Ripk3^-/-^*^58^*,Mlkl^-/-^*^58^ *and Tnfr1^-/-^*^59^ mouse lines have been described earlier. The mouse *Cyld* D215A mutation construct corresponds to the following genomic position (NC_000074.7) chr13: 89423506-89478574. *Cyld^D^*^215^*^A/+^* mouse genotyping primers: 5′-ATTATGATTTGAAACGATGAGG-3′ and 5′-CTGGACCAAAGATAAACTGAA-3′ amplified 627□bp DNA fragments for sequencing. *Cyld^+/-^* mice were generated by a CRISPR-Cas9 approach using the following sequence: sing-guide RNA (sgRNA) target 1, 5′- TACTGTCCTATACTCCCTTT-3′; sgRNA target 2, 5′- GGGACTTACAGCGAGTTCAT-3′. CYLD mice genotyping primers (CYLD-F: 5′- GGGACTTACAGCGAGTTCAT-3′ and CYLD-R: 5′- ATAGATCAGTGGTAGAGGGT-3′) amplified 778 bp wild-type, 349□bp knock-out DNA fragments. The *Lysm^cre+^* mice were purchased from Shanghai Model Organisms Center. For *Rosa26^CAG-LSL-EGFP-Cyld/+^* mice, the cDNA encoding *cag-loxp-stop-loxp-EGFP-Linker-Cyld* was generated and inserted into intron between exon1 and exon2 of the *Rosa26* locus. The Rosa26^CAG-LSL-EGFP-Cyld/+^ mice were bred with *LysM^cre+^* mice to generate *Rosa26^CAG-LSL-EGFP-Cyld/+^ LysM^cre+^* mice. Additional information is provided upon request. All mice utilized in this study have been backcrossed onto the C57BL/6 background for more than eight generations. Animal experiments were conducted in accordance with the guidelines of the Institutional Animal Care and Use Committee of the Institute of Nutrition and Health, Shanghai Institutes for Biological Sciences, University of Chinese Academy of Sciences.

For TLR4-dependent endotoxic shock, mice were injected intraperitoneally with LPS from E. coli 055:B5 LPS (L2880, Sigma) at 50 mg/kg. For TLR4-independent endotoxic shock^11^, mice were primed by intraperitoneal injection of low molecular weight Poly(I:C) (LMW, InvivoGen; 4 mg/kg) followed 6h later by intraperitoneal challenge with LPS (L2630, Sigma) at 5 mg/kg. Injection volume for all experiments was 200μl made with sterile PBS. For the experiment in the **Fig. 6n**, mice were injected intraperitoneally with HOIPIN-1 from MCE (HY-122881) at 3mg/kg followed 6h later by intraperitoneal challenge with LPS at 50mg/kg (L2880, Sigma). For imiquimod-induced psoriasis experiments, treated back was shaved prior to topical application of 62.5mg of MED SHINE (HGS045006C) cream for six consecutive days. Lesions were evaluated histologically 24 hours after the last application. Histologic images of skin were evaluated for epidermal hyperplasia (scored 1–4), crusting and ulceration (scored 1–4), and dermatitis and panniculitis (scored 1–4), with greater scores indicating worse disease. Total clinical score was calculated by adding these individual scores.

### Western blotting and immunoprecipitation

Cells and tissues were lysed in RIPA lysis buffer containing 50□mM Tris (pH□=□8.0), 1□mM EDTA, 150□mM NaCl, 0.1% SDS, 1% Triton X-100, 0.2% sodium deoxycholate, 50□mM NaF, 1× proteinase inhibitor cocktail (Roche, 5056489001), 2□mM PMSF, and 5□mM NEM. The lysates were cleared by centrifugation for 30□min at 12,000□×□g, quantified by BCA kit (P0010S, Beyotime), and then mixed with SDS sample buffer and boiled at 96□°C for 10□min. For TUBE pull-down, cells were stimulated with Pam3csk4 or LPS at indicated time, and lysed with TUBE lysis buffer (100□mM Tris-HCl, pH 8.0, 0.15□M NaCl, 5□mM EDTA, 1% NP-40, 0.5% Triton-X 100, 2□mM NEM, 50□µM PR619, 2□mM o-PA, protein inhibitor cocktail, 5□mM PMSF and 150nM flag-TUBE) at 4 °C for 30□min. The lysates were cleared by centrifugation and then diluted ten-fold in dilution buffer (100□mM Tris-HCl, pH 8.0, 0.15□M NaCl, 5□mM EDTA, 2□mM NEM, 50□µM PR619, 2□mM o-PA, protein inhibitor cocktail, 5□mM PMSF, and 150nM flag-TUBE). Flag-TUBEs (UM-0401-1000, LifeSensors) were pulled down with anti-flag beads overnight at 4□°C, and then washed with cold wash buffer (100□mM Tris-HCl, pH 8.0, 0.15□M NaCl, 5□mM EDTA, 0.05% NP-40). Finally, 20□µl of 5× SDS reducing loading sample buffer was added to the resin, and the sample was boiled at 96□°C for 10□min.

For polyubiquitination assays, cells were washed twice with ice-cold 1× PBS and subsequently lysed in NP-40 buffer (0.5% NP-40, pH 7.5, 120□mM NaCl, 10% glycerol, 30□mM Tris, 1□mM EDTA, 2□mM KCl, 50□mM NaF, 5□mM sodium pyrophosphate, 10□mM β-glycerophosphate, 2□mM PMSF, 10□mM NEM, and 1x proteinase inhibitor cocktail) at 4□°C for 30min with rotation. The lysates were cleared by centrifugation and then quantified by BCA kit. The normalized lysates were subsequently denatured in reducing 5×SDS loading sample buffer at 100□°C for 10□min. The normalized supernatants were mixed with FLAG-tagged beads (Sigma, A2220) at 4°C overnight. The beads were washed with ice-cold NP-40 buffer at least three times. Proteins were eluted with 2×SDS loading sample buffer and boiled at 96□°C for 10□min.

The samples were separated using SDS-PAGE, transferred to PVDF membrane (Millipore) with 110□v for 2□h. The proteins were detected by using a chemiluminescent substrate (34580, Pierce).

The following antibodies were used for western blotting and immunoprecipitation experiments: Caspase8 (1:2000, Enzo Life Science, ALX-804-447-C100), MLKL (1:2000, Abgent, ap14272b), CYLD (1:2000, Cell Signaling Technology, 8462S), β-actin (1:10000, Sigma, A3854), Caspase1 (1:2000, Adipogen, AG-20B-0042), GSDMD (1:2000, Cell Signaling Technology, 97558S), Cleaved-GSDMD(1:2000, Cell Signaling Technology, 10137SS), NLRP3 (1:2000, Adipogen, AG-20B-0014-C100), Caspase11 (1:2000, Cell Signaling Technology, 14340S), p-IKKα/β (1:2000, Cell Signaling Technology, 2697S), IKKα (1:2000, Cell Signaling Technology, 2682S), IKKβ(1:2000, Cell Signaling Technology, 8943S), IkBα (1:2000, Cell Signaling Technology, 9242S), p-IkBα (1:1000, Cell Signaling Technology, 9246S), p-P65 (1:2000, Cell Signaling Technology, 3033S), P65 (1:2000, Cell Signaling Technology, 4764S), P65(1:2000, Cell Signaling Technology, 8242S), RIPK3 (1:5000, Prosci, 2283), FADD (1:1000, Abcam, ab124812), GAPDH (1:20,000, Sigma, G9545), TBP (1:2000, Cell Signaling Technology, 44059S), HA-Tag (1:2000, Cell Signaling Technology, 3724S), Flag-Tag (1:2000, Sigma, A8592-1MG), Myc-Tag (1:2000, Cell Signaling Technology, 2276S), GFP (1:1000, Cell Signaling Technology, 2956S), HSP90 (1:5000, Cell Signaling Technology, 4874S).

### Transfection of Hela and 293T cells

Hela and 293T were transfected using Lipofectamine 2000 (11668019, Invitrogen). Transfections were according to manufacturer’s instructions. For 1 well of a 6-well plate, 1μg of DNA was diluted in 100μl opti-MEM medium (31985070, Gibico) before adding 100μl opti-MEM medium containing 2μl Lipofectamine.

### ELISA of cytokines

Mouse serum and cell supernatants were assayed using TNF-α (eBioscience, 88-7324-88), IL-1β (eBioscience, 88-7013-88), IL-18 (eBioscience, BMS618/3) and IL-6 (eBioscience, 88-7064-88) ELISA kits according to manufacturer’s instructions.

### Plasmids

HA-CYLD, HA-IKKα, HA-IKKβ, HA-IκBα HA-importin α1, HA-importin α3 and HA-importin α4 were generated by cloning mouse *CYLD* cDNA, *IKK*α cDNA, *IKK*β cDNA, *I*κ*B*α cDNA, *importin* α*1* cDNA, *importin* α*3* cDNA and *importin* α*4* cDNA into the pcDNA3.0-3×HA vectors. Mouse *CYLD* cDNA was cloned into pcDNA3.0-Flag-eGFP vectors and pcDNA3.0-Myc vectors. Flag-P65 was generated by cloning mouse *P65* cDNA into the pCDH-3xFlag vectors and pCDH-3xFlag-mCherry vectors. Myc-HOIP, Myc-HOIL-1 and Myc-Sharpin were generated by cloning mouse *HOIP* cDNA, *HOIL-1* cDNA and *Sharpin* cDNA into the pcDNA3.0-Myc vectors.

Point mutations were generated by site-directed mutagenesis. The Flag-P65 mutants ΔRHD, ΔIPT, ΔPHA, K62R, K122&123R, K195R, K218R, K301R, K303R, K301&303R, K310R, K314&315R and K425R were generated using Flag-P65 as the template. The 3×HA-CYLD mutants ΔPhos, ΔCG3, ΔDUB were generated suing 3×HA-CYLD as the template. The Myc-CYLD mutans C600A and C600S were generated using Myc-CYLD as the template. The Myc-HOIP mutants C879A and C910A were generated using Myc-HOIP as the template.

The following plasmids were gifts from Hao Ying: HA-Ubiquitin WT, K6, K11, K27, K29, K33, K48, K63, K48R, K63R. HA-Ubiquitin M1 and M1A were generated by site-directed mutagenesis using HA-Ubiquitin WT as the template.

### CRISPR-Cas9-mediated depletion of CYLD

For the depletion of CYLD, single guide RNA (sgRNA) targeting *CYLD* was cloned by annealing two DNA oligos (forward, 5’-GTGGTCAAGGTTTCACTGAC-3’; reverse, 5’-GTCAGTGAAACCTTGACCAC-3’) and ligating into lentiCRISPER v2 (52961, Addgene) plasmids. 293T and Hela cells were infected with lentiCRISPER v2-sgCYLD vectors and selected in the presence of puromycin.

### Luciferase reporter assay

Hela and 293T cells were plated in 12-well plates and transfected with pRL-TK and plasmids encoding the NF-κB luciferase reporter, together with different plasmids following: Flag-P65, Flag-P65-K62R, Flag-P65-K122&123R, Flag-P65-K195R, Flag-P65-K218R, Flag-P65-K301&303R, Flag-P65-K310R, Flag-P65-K314&315R, Flag-P65-K425R, Flag-NEMO, Flag-TRAF2, Myc-CYLD, Myc-CYLD-C600A,

Myc-CYLD-C600S or empty vectors. At 24h after transfection, cells were lysed in native lysis buffer and luciferase activity was assayed using the Dual-Luciferase Reporter Assay (E1910, Promega) according to the manufacturer’s instructions.

### Quantitative real-time PCR

Total cellular or skin RNA was extracted using Trizol reagent (9108, Takara), and cDNA was synthesized by using the Reverse Transcriptase kit (RR037A, Takara). Real-Time PCR was performed by using specific primers and SYBR Green ROX Mix (RR420A, Takara). β-actin was used as the internal quantitative control.

Primer used are as follows:

mIL-6-F: 5’- GGGAAATCGTGGAAATGAGA-3’

mIL-6-R: 5’- CCAGTTTGGTAGCATCCATCA-3’

mNLRP3-F: 5’-ATTACCCGCCCGAGAAAGG-3’

mNLRP3-R: 5’-TCGCAGCAAAGATCCACACAG-3’

mCaspase11-F: 5’- ACAAACACCCTGACAAACCAC-3’

mCaspase11-R: 5’- CACTGCGTTCAGCATTGTTAAA-3’

### Cell Culture

Mouse embryonic fibroblasts (MEFs) were cultured in Dulbecco’s Modified Eagle’s Medium (SH30243.LS, Hyclone) supplemented with 10% fetal bovine serum (SH30084.03, Hyclone), 1% penicillin (100□IU/ml)/streptomycin (100□μg/ml) and 2□mM L-glutamine under 5% CO2 at 37□°C. Primary MEFs were isolated from E11.5 littermate embryos. The head and visceral tissues were dissected and remaining bodies were incubated with 4□ml trypsin/EDTA solution (25200072, Invitrogen) per embryo at 37□°C for 1□h. After trypsinization, an equal amount of medium was mixed and pipetted up and down several times. The primary MEFs were cultured in high-glucose DMEM containing 15% FBS. For immortalization, MEFs were transfected with SV40 small+large T antigen-expressing plasmid (22298, Addgene) using Lipofectamine 2000 (11668019, Invitrogen) according to manufacturer’s instructions.

For culturing bone marrow-derived macrophages, bone marrow progenitors harvested from femurs and tibias of age- and sex-matched 8- to 12-week-old mice and differentiated into macrophages over 7 days in RPMI 1640 with M-CSF (conditioned medium from L929 cell line) and 10% fetal bovine serum, plus 100U/mL penicillin and 100μg/mL streptomycin.

Peritoneal macrophages were cultured in RPMI 1640 with 10% fetal bovine serum, plus 100U/mL penicillin and 100μg/mL streptomycin. Peritoneal macrophages were isolated from 6- to 8-weeks-old C57BL/6J mice. 3 days after intraperitoneal injection of 2 mL of 3% thioglycollate medium (BD, Cat#: 211716), peritoneal macrophages were isolated and cultured in RPMI 1640 medium supplemented with 10% FBS (HyClone, Cat#: SH30084.03), 100 U/mL penicillin, and 100 mg/mL streptomycin for use.

### Chloroform methanol purification of secreted CP25

Mouse BMDMs were stimulated with LPS (100ng/ml, O55:B5, Sigma-Aldrich) in RPMI 1640 without serum for indicated times and the cell culture medium (supernatant) was collected and filtered using a membrane filter unit (SLGPR33RS, Millipore). Chloroform methanol purification of proteins from the filtered supernatant was performed and dried proteins were eluted with 1× SDS loading sample buffer and boiled at 96□°C for 10□min.

### Statistical analysis

Data presented in this article are representative results of at least three independent experiments. A log-rank (Mantel-Cox) test was used to compare mouse survival. The statistical significance of data was evaluated by Student’s t test. The correlation analysis was determined by linear regression. The statistical calculations were performed with GraphPad Prism software.

## Supplementary Information

Supplementary information includes six figures.

## Author Contributions

H.B.Z. conceived the project. J.L.L., M.L. and H.B.Z designed the study; J.L.L. performed the experiments and data analyses with assistance from M.Y.X., H.L., L.X.W., X.M.L., F.L., X.M.Z., Y.J.O., X.X.W., Y.Y.W., Y.Y.X. and H.W.Z; X.Q. assisted in cell death analysis; Y.Z. and Y.C.T. contributed to the histopathological analysis; H.L. and Y.L contributed to critical discussion. Y.L. and H.L. provided essential resources and intellectual input during the experimental studies. J.L.L., M.L. and H.B.Z assembled figure panels and wrote the paper. H.B.Z supervised the project.

## Acknowledgements

We thank Dr. Guangxun Meng for providing *Caspase1/11^-/-^* mice and Dr. Hui Xiao for providing *Trif^-/-^* mice. This work was supported by grants from National Key Research and Development Program of China (2022YFA0807300), the Strategic Priority Research Program of Chinese Academy of Sciences (XDA26040306), National Natural Science Foundation of China (32270803, 31970688, 82272181). Shanghai Excellent Academic/Technical Leader Program (22XD1404500) and Shanghai Science and Technology Commission (23141902800). We also thank support from Shanghai Municipal Science and Technology Major Project, Shanghai Frontiers Science Center of Cellular Homeostasis and Human Diseases, and GuangCi Professorship Program of Ruijin Hospital Shanghai Jiao Tong University School of Medicine.

## Supplementary Figure Legends

**Figure S1. Generation and characterization of *Cyld^D^*^215^*^A/D^*^215^*^A^* mice.**

**a-c**, Western blot of BMDMs stimulated with LPS (100ng/ml, O55:B5) with or without Chloroquine (CQ) **(a)**, MG132 **(b)** and zVAD **(c)** treated for indicated times. **d**, Western blot of WT, *Mlkl^-/-^* and *Casp8^-/-^Mlkl^-/-^* BMDMs stimulated with LPS (100ng/ml, O55:B5) treated for indicated times.

**e**, Aspartic acid (GAT) was mutated to Alanine (GCT) at the 215 position in CYLD. The mutation was confirmed by sequencing.

**f**, Western blot of lung, thymus and heart from WT and *Cyld^D^*^215^*^A/D^*^215^*^A^* mice.

**g**, Western blot of lung, bone marrow, spleen and liver from WT and *Cyld^D^*^215^*^A/D^*^215^*^A^* mice injected intraperitoneally with 25mg/kg of LPS (O55:B5).

**Figure S2. CYLD regulates of P65 activation and inflammatory cytokines release.**

**a-b**, WT and *Cyld^D^*^215^*^A/D^*^215^*^A^***(a)**; WT and *Trif^-/-^* **(b)** BMDMs were stimulated for 1h with indicated TLR agonists, followed by immunoblot analysis.

**c**, Western blot of WT and *Cyld^-/-^* PMs stimulated with Pam3cks4 for indicated times. **d**, WT and *Cyld^-/-^* PMs stimulated with Pam3cks4 for indicated times. Then separate the nucleus and cytoplasm by Nuclear and Cytoplasmic Protein Extraction Kit and perform subsequent Western blot.

**e-f**, Plots show TNFα and IL-6 secretion of WT and *Cyld^-/-^* PMs treated with Pma3cks4 (1μg/ml) for indicated times.

**g**, Western blot of *Tnfr1^+/+^* and *Tnfr1^-/-^* BMDMs.

**h**, Western blot of WT and *Trif^-/-^* BMDMs stimulated with TNFα and FasL.

**i**, Western blot of immortalized MEFs with indicated genotypes treated with TNFα.

**j-k**, WT and *Cyld^D^*^215^*^A/D^*^215^*^A^* BMDMs were stimulated with FasL **(j)** and TNFα **(k)**, followed by immunoblot analysis.

P values calculated using t-test **(e and f)**. ^D^P<0.05, ^DD^P<0.01, ^DDD^P<0.001.

**Figure S3. CYLD deubiquitinates M1-linked ubiquitination of P65.**

**a-b**, J774 cells **(a)** and PM cells **(b)** were stimulated with LPS (100ng/ml, O55:B5) and Pmascsk4 (1μg/ml) correspondingly for indicated times. Cell lysates were subjected to immunoprecipitation using TUBE antibody and Flag-beads. TUBE-Ub samples and cell lysates (input) were analyzed using Western blot.

**c**, Schematic showing the different HA-tagged plasmids of ubiquitination mutations. **d**, 293T cells were transfected with Flag-P65, Myc-CYLD and HA-tagged ubiquitin plasmids with the indicated combinations for 24h. The cell lysates were subjected to immunoprecipitation with anti-Flag beads and immunoblotted with the indicated antibodies.

**e-f**, 293T cells were transfected with Flag-P65, Myc-CYLD, Myc-CYLD-C600S, HA-WT **(e)** and HA-M1 **(f)** plasmids with the indicated combinations for 24h. The cell lysates were subjected to immunoprecipitation with anti-Flag beads and immunoblotted with the indicated antibodies.

**g-h**, 293T **(g)** and Hela **(h)** cells stably expressing control or *Cyld* small-guide RNA. And then cells were transfected with Flag-P65 and HA-M1 plasmids as indicated for 24h. The cell lysates were subjected to immunoprecipitation with anti-Flag beads and immunoblotted with the indicated antibodies.

**Figure S4. CYLD coordinating with Lubac complex regulates M1-linked ubiquitination of P65 on K301/303 residues.**

**a**, 293T cells were transfected with Flag-P65, Myc-HOIP, Myc-HOIL-1 and Myc-Sharpin, and plasmids with the indicated combinations. The cell lysates were subjected to immunoprecipitation with anti-Flag beads and immunoblotted with the indicated antibodies.

**b**, Prediction of mP65 ubiquitination sites.

**c**, Hela cells were transfected with HA-WT, Flag-P65 (and its mutants) and plasmids with the indicated combinations. The cell lysates were subjected to immunoprecipitation with anti-Flag beads and immunoblotted with the indicated antibodies.

**d**, 293T cells were transfected with pRL-TK, NF-κB luciferase reporter, empty vector, Flag-P65 and its mutans. Then NF-κB activity was assayed.

**e**, Hela cells were transfected with HA-WT, Flag-P65 (and its mutants) and plasmids with the indicated combinations. The cell lysates were subjected to immunoprecipitation with anti-Flag beads and immunoblotted with the indicated antibodies.

**f**, Hela cells were transfected with Flag-P65 and its mutant Flga-P65-DM for 24h. Then cells were stimulated with mTNFα for indicated times. The cytoplasmic and nuclear extracts perform subsequent Western blot.

P values calculated using t-test **(d)**. ^D^P<0.05, ^DD^P<0.01, ^DDD^P<0.001.

**Figure S5. CP25 secretion is independent of unconventional secretory pathways.**

**a**, Western blot of cytoplasmic and nuclear extracts from BMDMs stimulated with LPS (100ng/ml, O55:B5) for indicated times.

**b**, BMDMs were pretreated with 0μm, 10μm, and 20μm GW4869 for 2 hours, then stimulated with LPS (100ng/ml, O55:B5) for 5 hours followed by determination of CP25 secretion.

**c**, Schematic showing the Lysm-cre+ induced expression of EGFP-CYLD in macrophages in Rosa26CAG-LSL-EGFP-Cyld/+ mice.

**d**, Summary model for mechanisms by which the ubiquitination of P65 mediated by CYLD that cleaved by Caspase8.

P values calculated using t-test **(d)**. ^D^P<0.05, ^DD^P<0.01, ^DDD^P<0.001.

