## Supplementary figures and images for "Caspase-8-mediated CYLD Cleavage boosts LPS-induced Endotoxic Shock"

### Supplemental Figure 1

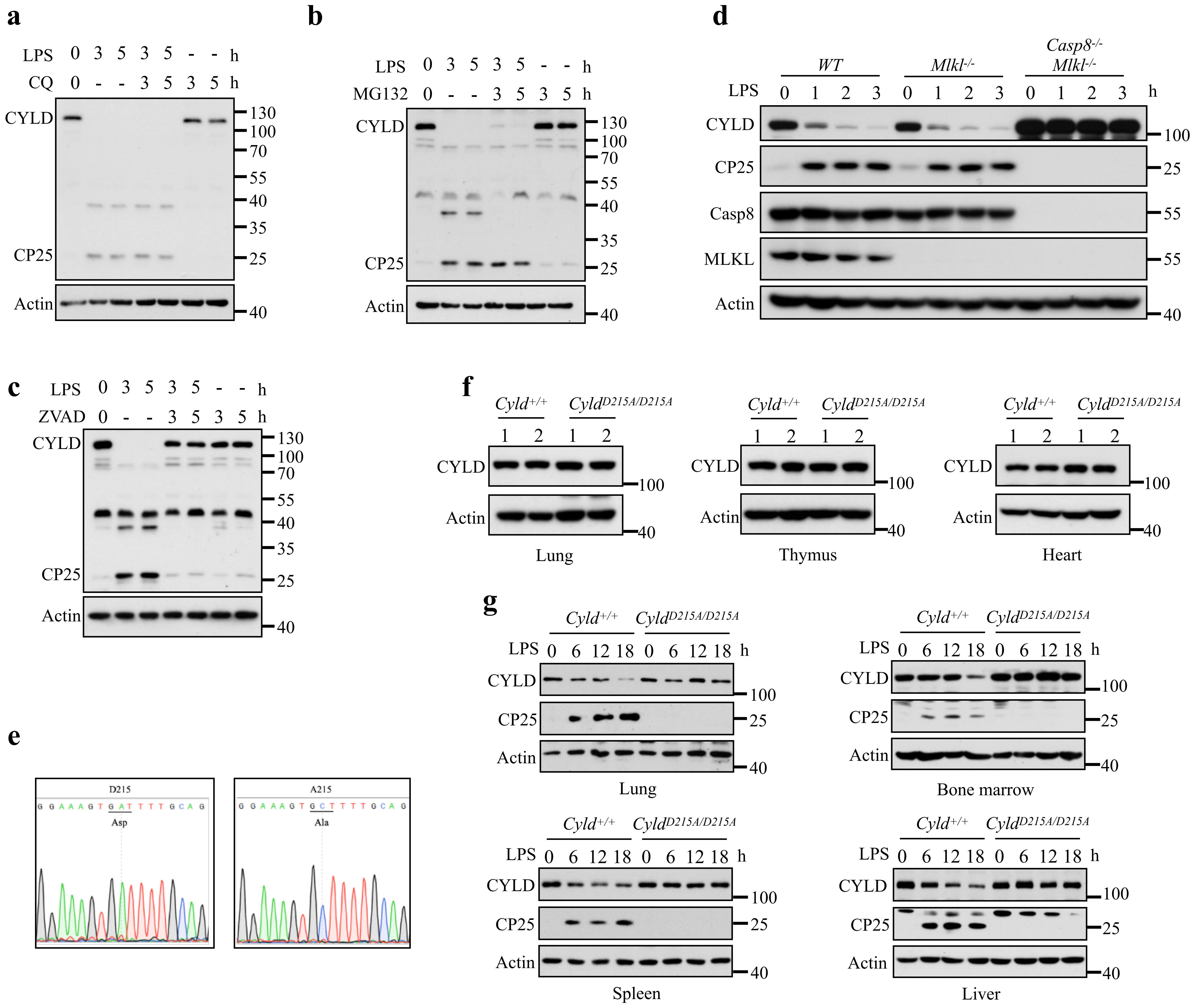

### Supplemental Figure 2

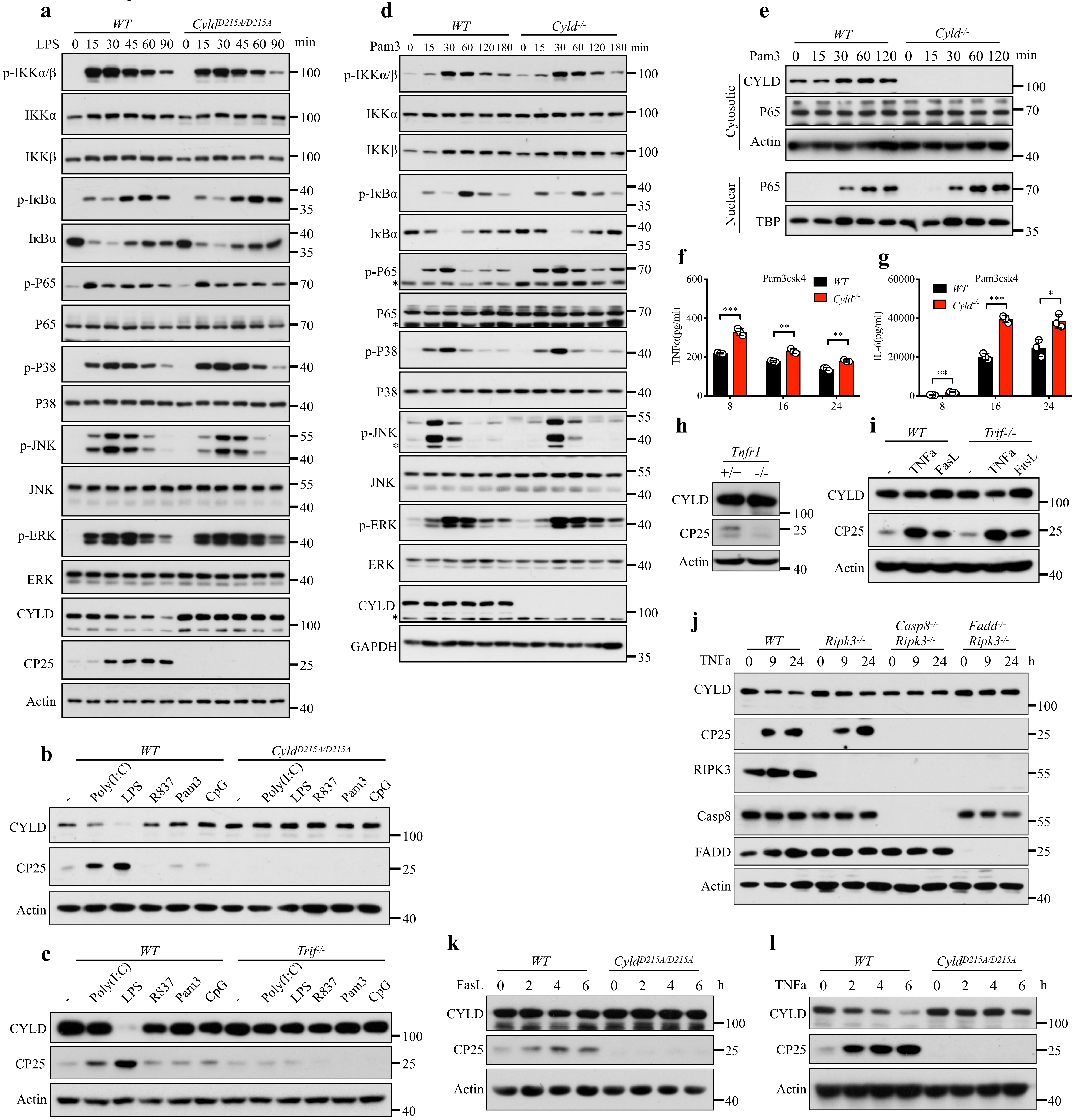

### Supplemental Figure 3

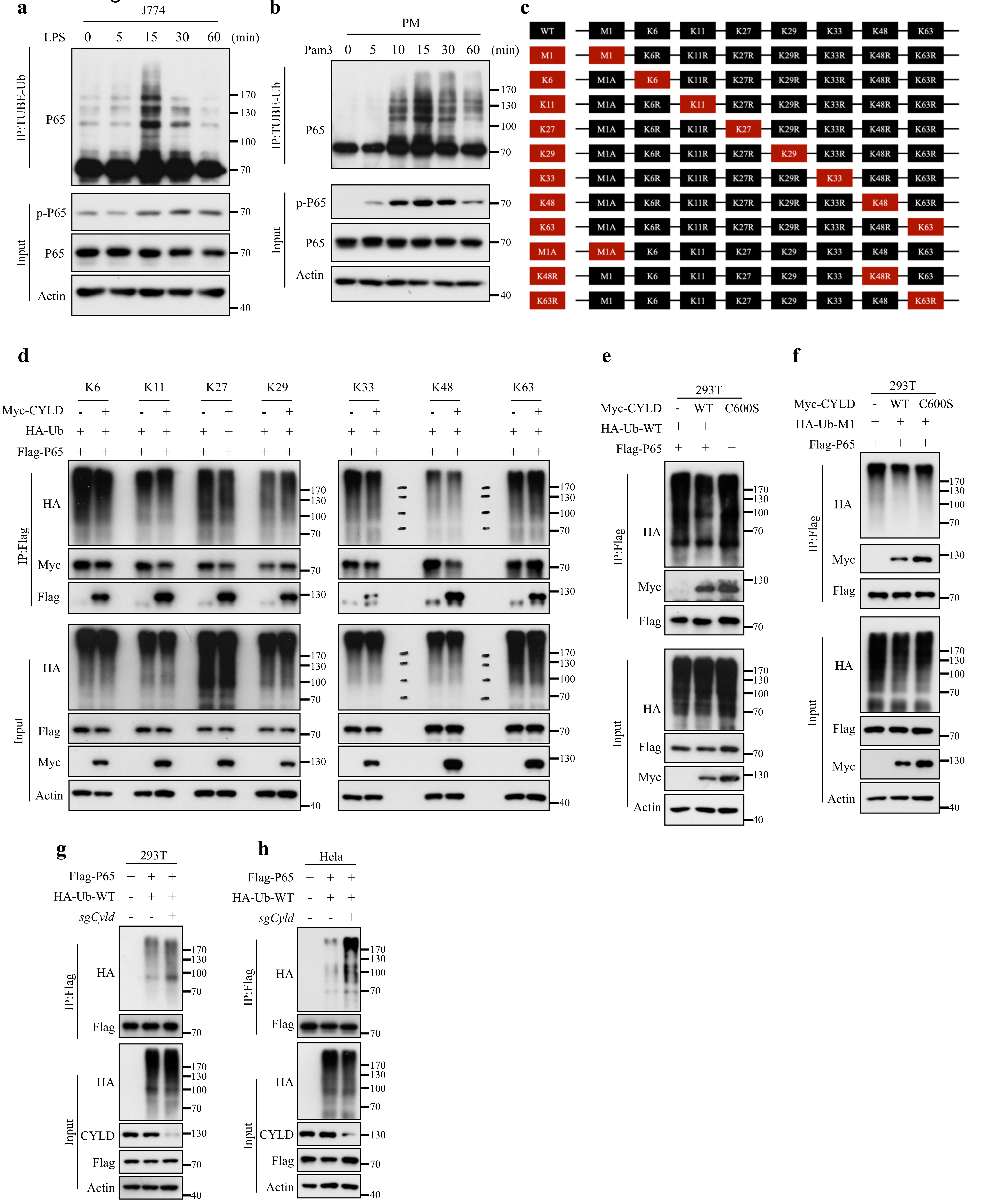

### Supplemental Figure 5

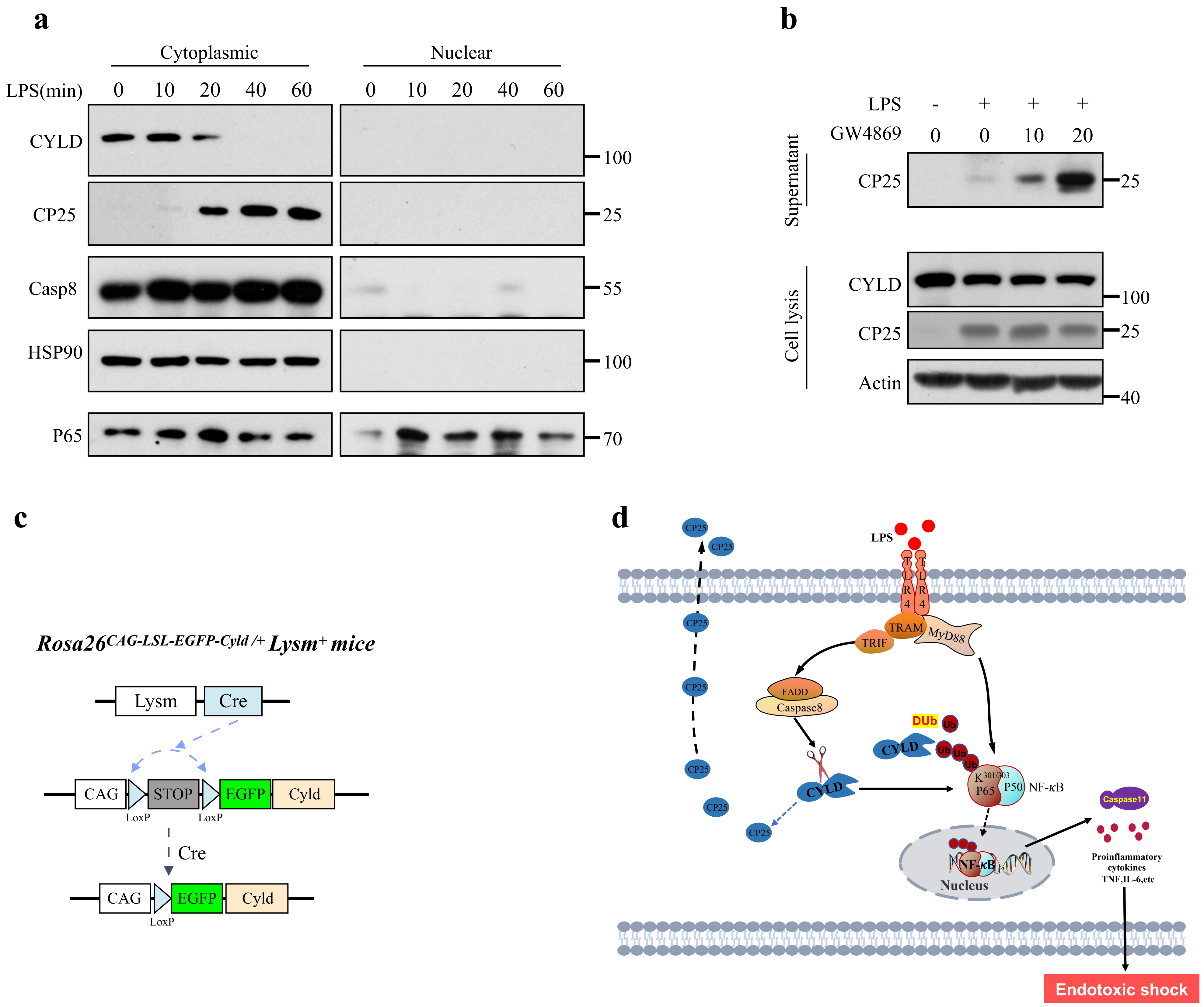
