## Supplemental Figure 4 for "Caspase-8-mediated CYLD Cleavage boosts LPS-induced Endotoxic Shock"

### Figure S4

**a**

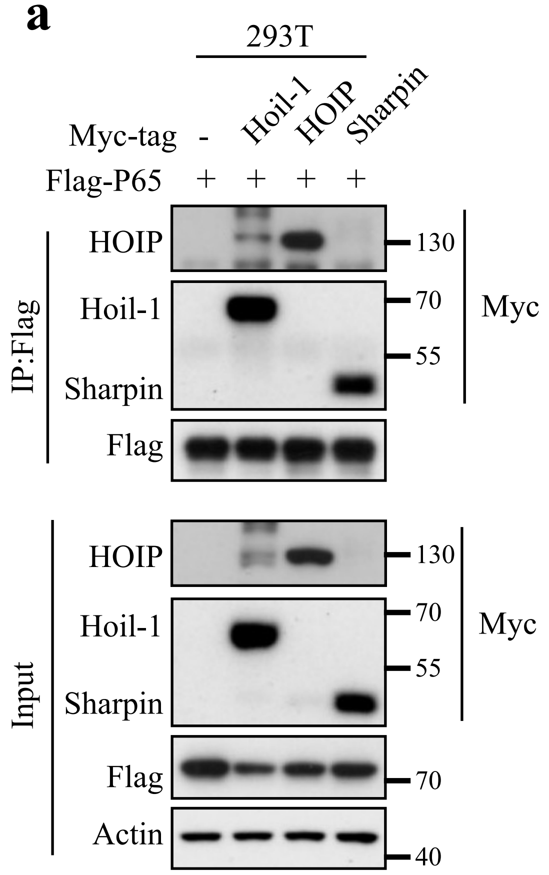


**b**

Predicted mP65 ubiquitination sites

| Peptide | Position | Score |
|---|---|---|
| TKTHPTIKINGYTGP | 62 | 0.8 |
| NLGIQCVKKRDLEQA | 122 | 0.4 |
| LGIQCVKKRDLEQAI | 123 | 1.56 |
| APNTAELKICRVNRN | 195 | 1.94 |
| EIFLLCDKVQKEDIE | 218 | 1.28 |
| DRHRIEEKRKRTYET | 301 | 0.54 |
| HRIEEKRKRTYETFK | 303 | 0.69 |
| KRTYETFKSIMKKSP | 310 | 1.33 |
| ETFKSIMKKSPFNGP | 314 | 0.66 |
| TFKSIMKKSPFNGPT | 315 | 1.32 |
| SLSAPVPKSTQAGEG | 425 | 1.97 |

**c**

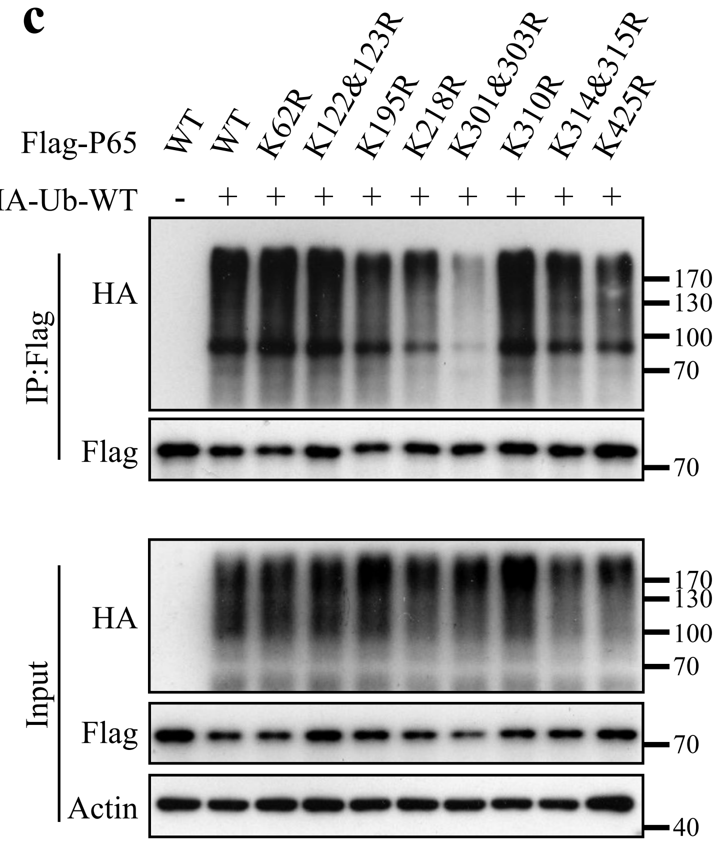


**d**

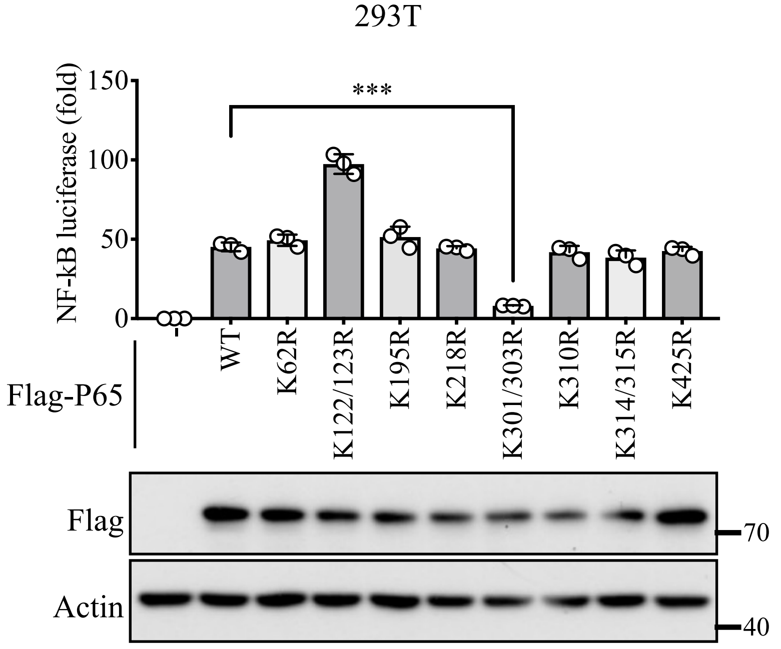


**e**

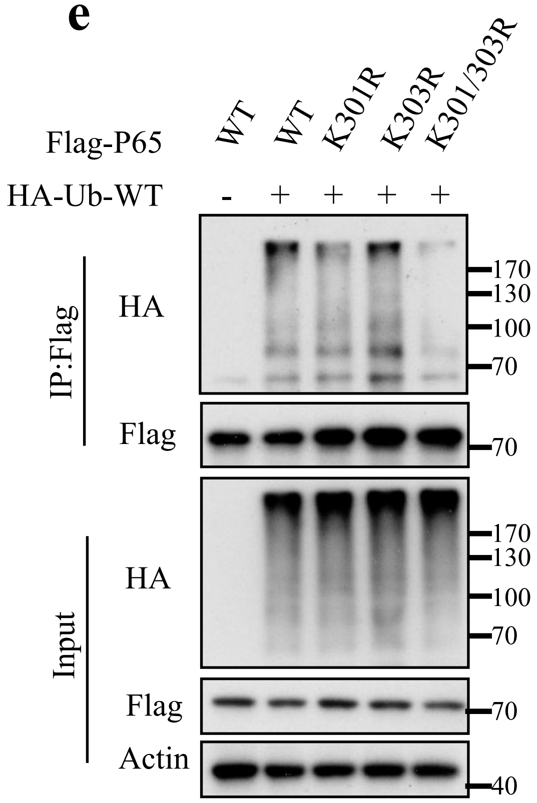


**f**

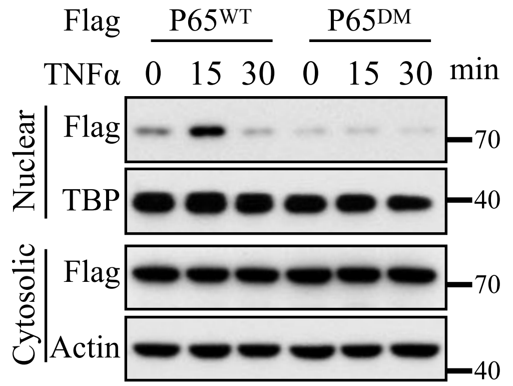
